# Synthetic Circuits Based on Split CAS9 to Detect Cellular Events

**DOI:** 10.1101/2023.03.16.533022

**Authors:** Alicja Przybyszewska-Podstawka, Jakub Czapiński, Joanna Kałafut, Adolfo Rivero-Müller

## Abstract

Synthetic biology involves the generation of logic circuits to create or control biological functions and behaviors by engineering interconnected genetic elements such as promoters, repressors, and transcriptional activators. CRISPR discovery, and its adaptations to mammalian cells, has made it the tool of choice in molecular biology and revolutionized genome engineering in biomedical sciences. Here, we describe an adaptation of a split Cas9 to generate synthetic logic gates to sense biological events. As proof-of-concept, the complementing halves of split Cas9 were placed under different promoters, one unique to cancer cells of epithelial origin (_*p*_*hCEA*) and one universal promoter (_*p*_*CMV*). We used self-assembling inteins to reunite the halves when co-expressed. Only cancer cells with epithelial origin expressed both halves and activated a reporter becoming green fluorescent. We then investigated whether we could apply this system to the detection of biological processes such as epithelial to mesenchymal transition (EMT). We designed another logic gate where one halve is expressed only by cancer cells of epithelial origin, while the other is activated during EMT – under the control of TWIST1. Indeed, cells undergoing EMT were detected by the activation of the reporter. Finally, the split-Cas9 logic gate was applied as a sensor to detect cell-cell fusion events in multiple cell lines. Each cell type expressed only one halve of split Cas9, and only induction of fusion resulted in the appearance of multinucleated syncytia and the expression of the reporter system. The simplicity and flexibility of the split Cas9 system reported here can be integrated to many other cellular processes, not only as a sensor but as an actuator.

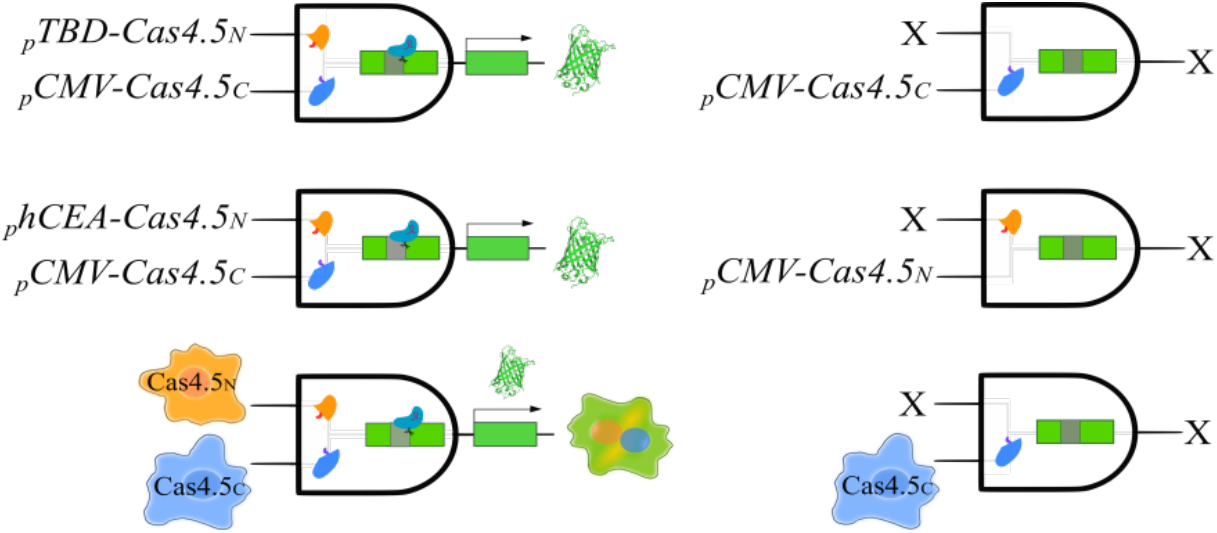

## INTRODUCTION

In synthetic biology, most of the logic control developed so far employ transcriptional regulators e.g., transcriptional activators or repressors to regulate gene expression^1–5^ to either increase reaction rates in metabolic pathways or respond to new input signals. While these genetic circuits work very predictably in prokaryotes, in eukaryotes they often require the use of genetic elements from prokaryotes or simple eukaryotes.

The domestication of CRISPR systems, the prokaryotic “immune system” that confers resistance and memory to phageal genetic elements, to eukaryotic cells has opened a new level of genetic engineering. CRISPR/Cas9 is composed by Cas9 endonuclease, responsible for executing the desired genetic modifications, and two small RNA molecules named CRISPR RNA (crRNA) and trans-activating CRISPR RNA (tracrRNA). While crRNA contains targeting sequence, the tracrRNA acts as a scaffold for Cas9 endonuclease^6^. In order to simplify the system, the crRNA and tracrRNA have been fused into a single guide-RNA (gRNA)^7–9^. Cas9/gRNA generates double strand DNA breaks which can be repaired by either nonhomologous end-joining pathway (NHEJ), causing errors such as insertion/deletions (indels), or by homology-directed repair (HDR), which can be used to introduce specific DNA sequence changes. Such simplicity of CRISPR systems has made them the most adaptable system to manipulate endogenous genetic information, from gene deletion (knockout)^10–13^, gene insertion (knock-ins)^14–16^, to gene regulation (CRISPRi and CRISPR-TA)^17^, and genome editing^18–20^.

The Cas9 system has also been integrated into genetic logic gates whereas the input is provided by gRNAs that respond to a particular stimulus, such as the presence of a specific chemical^21,22^ or the activation of a specific signaling pathway, or regulate multiplexed circuits to control transcriptional regulation (activation/inhibition) by modified Cas9 forms (CRISPRa and CRISPRi, respectively)^23–26^.

Yet, CRISPR/Cas9 has been rarely used as a sensor for biological events i.e. for identification of cancer cells^27^ or to control a complex phenotype such as cell fate^28^. In these cases, Cas9 and the gRNA have been placed under specific promoters, each becoming one input to the AND logic gate. A split Cas9 system has previously been reported although not applied into logic circuits^29^, although it has been used to generate drug- and light-inducible systems by fusing halves of Cas9 to hetrodimerizers^30,31^.

Here, we present another use of CRISPR/Cas9 by creating a synthetic (AND) logic gate using a split Cas9 (named Cas4.5_N_ and Cas4,5_C_ -as they equal 9) that need to be co-expressed (input 1 and 2) to activate a reporter, and leads to a specific output (outcome). While studying cellular events can be detected by biochemical assays or biosensors, there are very few examples on the use of synthetic logic gates to record cellular events in eukaryotic cells.

Moreover, the presented 2-input system results in an active actuator that can be applied to many other biological outcomes. We show that detection of cell origin, phenotypical transition or cell-cell fusion can be specifically detected by split Cas9 logic gates.

## EXPERIMENTAL SECTION

### Materials and Reagents

Plasmid pSpCas9(BB)-2A-Puro (PX459) V2.0 was a gift from Feng Zhang (Addgene plasmid #62988^32^), pU6-sgGFP-NT1 was a gift from Stanley Qi & Jonathan Weissman (Addgene plasmid #46914; http://n2t.net/addgene:46914; RRID:Addgene_46914)^8^), pCAG-EGxxFP-*Cetn1* was a gift from Masahito Ikawa (Addgene plasmid #50717; http://n2t.net/addgene:50717; RRID:Addgene_50717^33^), AmCyan-P2A-mCherry was previously created by us^34^ (Addgene plasmid #45350; http://n2t.net/addgene:45350; RRID:Addgene_45350^34^). The TWIST1 binding domain (*TBD*) promoter was created previously by us^35^, DnaE intein plasmids were a gift from Hideo Iwai (Addgene plasmid #34549; http://n2t.net/addgene:34549; RRID:Addgene_34549^36^ and #15335; http://n2t.net/addgene:15335; RRID:Addgene_15335^37^). Cell lines HEK-293, HSF, H1299, H2170, SW480, A375 and HeLa, were obtained from ATCC; H2170 stable cell line with vimentin reporter (VRCs) was created by us^38^. KOD-Xtreme hot-start DNA polymerase (Merck Millipore), DreamTaq™ Green PCR Master Mix (ThermoFisher Scientific), endonuclease *DpnI* and *BbsI* (Thermo Fisher Scientific), T7 Ligase (Thermo Fisher Scientific) and PNK (Thermo Fisher Scientific), Polyethylene Glycol Hybri-Max 1450 (PEG) (Sigma-Aldrich), TGFβ (human TGFβ1, Biorbyt, Cambridge, United Kingdom), Gibson Assembly^®^ Master Mix (NEB), Ampicillin (BRAND), DNA Clean & Concentrator and Zyppy Plasmid Kits (Zymoresearch), Hoechst 33342 (Cayman), Dulbecco’s Modified Eagle Medium (DMEM/F-12), Penicillin and Streptomycin (Sigma Aldrich), fetal bovine serum (PromoCell) and geneticin G418 (ThermoFisher Scientific). All primers were bought from Genomed (Warsaw, Poland).

### PCR

All primers were designed in SnapGene Viewer and purchased from Genomed (Warsaw, Poland), Syncytin1 was recoded based on the amino acid sequence from the The Human Protein Atlas and ordered as a gBlock (ThermoFisher). PCR reactions were performed using a high-fidelity polymerase (KOD-Xtreme, Merck Millipore) according to manufacturer protocol. All products used for molecular cloning were then digested with fast digest *DpnI* restriction endonuclease (Thermo Scientific) to get rid of the methylated template plasmid. PCR products were column-purified before cloning using DNA Clean & Concentrator (Zymo research).

### Molecular cloning

All vectors were created using Gibson Assembly^39^ according to manufacturer protocol (NEB). In brief, inserts and vectors were amplified by PCR, or digested using restriction enzymes in selected cases, followed by gel-purification and concentrations measured. Appropriate amounts were used for Gibson reaction (ratio insert:vector 3:1). Then, electroporation with 2ul of the reaction was performed by using prepared electrocompetent DH10B *E*.*coli* bacteria, and after 45 minutes, bacteria were plated onto agar plates for selection with an antibiotic. A screening colony was taken using PCR, and the sequence was verified. The EGxxFP-*Cetn1* plasmid was modified by inserting the Puro-T2A cassette from pU6-sgGFP-NT1 (Addgene plasmid #46914) by recombination resulting in the Puro-T2A-EgxxFP plasmid. Cas4.5_N_ was amplified by PCR from Cas9 plasmid and then fused at its *C*-terminus to DnaE_N_ intein, while Cas4.5_C_ was fused at its *N*-terminus to DnaE_C_ intein. In the next step, the CMV general promotor (*pCMV*) in Cas4.5N-DnaE_N_ was replaced by *hCEA* or the TWIST1 binding domain *(TBD)* specific promoters/genetic elements. Resulting vectors: _p_hCEA-Cas4.5_N_-DnaE_N_, TBD-Cas4.5_N_-DnaE_N_, TBD-mCherry and _p_CMV-Cas4.5_C_-DnaE_C_. The gRNAs against EGxxFP-*Cetn1* was cloned into both the Cas9 and the Cas4.5_C_ plasmids, following the Zhang’s lab protocol^40^ using T7 ligase and *BbsI* (both from ThermoFisher).

### Cell culture and transfection

Human embryonic kidney 293 (HEK293) cells, human normal colon epithelial cells (CCD 841 CoTr), lung cancer cell lines (H1299, H2170, H2170 VRCs), malignant melanoma cell line (A375), cervical carcinoma (HeLa), and colon cancer cell line (SW480) cells were cultured under standard conditions in the incubator (37^°^C/5% CO_2_). Media DMEM/F12 (Sigma Aldrich) were supplemented with 10% Fetal Bovine Serum (PromoCell) and 1% Penicillin/Streptomycin solution. Transfections have been carried out using TurboFect reagent (Thermo Fisher), and Lipofectamine 3000^®^. Ratio DNA/reagent have been optimized for each cell line in order to obtain the best transfection efficiency.

One day before transfection H1299 and H2170 VRCs were seeded at density 5 × 10^4^ cells/well into a 24-well plate, and co-transfected with one half of Cas9 protein (_p_CMV-Cas4.5_C_) driven by general promoter, construct with EGxxFP reporter plasmid, *Cetn1* gRNA and second half of Cas9 (TBD-Cas4.5_N_) under a promoter activated during EMT – under the control of TWIST1, using Turbofect™ transfection reagent following the manufacturer’s protocol. Transfection with only one half of Cas9 protein was used as a negative control (Nc). Next day after transfection cells were exposed to TGFβ (10 ng/mL) for 72 h to evaluate the expression level of EMT-related gene *TWIST1*. All experiments were performed in at least triplicate.

Equally, 24h before transfection CCD 841 CoTr, A375, SW480, H1299 and HeLa cells were seeded at density 5 × 10^4^ cells/well into a 24-well plate, and co-transfected with one half of Cas9 protein (_p_CMV-Cas4.5_C_) driven by general promoter, construct with EGxxFP reporter plasmid, *Cetn1* gRNA and second half of Cas9 (_*p*_*hCEA-Cas4*.*5*_*N*_) under a epithelial cancer-specific promoter, using Lipofectamine 3000^®^ (CCD 841 CoTr, A375) or Turbofect™ (H1299, SW480 and HeLa) transfection reagents following the manufacturer’s protocol. Transfection with only one half of Cas9 protein was used as a negative control (Nc), and transfection contains mCherry was used as a transfection control. All experiments were performed in at least triplicate.

Cell lines HEK293, H1299, and H2170 were seeded at density 5 × 10^4^ cells/well into a 24-well plate 24h before transfection. Using Turbofect™ transfection reagent following the manufacturer’s protocol, cells were transfected with either one half Cas4.5_N_ with EGxxFP reporter plasmid, *Cetn1* gRNA, or the second halve _p_CMV-Cas4.5_C_ with Syncytin1 plasmid. Transfection with only one half of Cas9 protein was a negative control (NC). In PEG treatment, cells were transfected with either Cas4,5, gRNA and reporter plasmids, and next day after transfection cells were exposed to 50% solution of PEG for 3 min at RT to induce cell-cell fusion. Similarly, to create a stable cell lines HEK293, H1299, and H2170 cells were seeded at density 5 × 10^4^ cells/well into a 24-well plate one day before transfection. Using Turbofect™ transfection reagent following the manufacturer’s protocol, cells were transfected with each halves (_p_CMV-Cas4.5_N_; _p_CMV-Cas4.5_C_). After 48 hours, cells were selected with Puromycin (0.75-1 mg/ml) for over three weeks to generate stable lines. Then, stable cell lines were co-transfected – one half Cas4.5_N_ with EGxxFP reporter plasmid, *Cetn1* gRNA, and second halve _p_CMV-Cas4.5_C_ with Syncytin1 plasmid.

### Wound-healing assay

For the Wound-healing assay cell lines of H1299 and H2170 VRCs (vimentin reporter cells)^38^ were seeded into 12-well plates at a density 5 × 10^4^ cells/mL. One day later were co-transfected with one half of Cas9 protein (_p_CMV-Cas4.5_C_) driven by general promoter, construct with EGxxFP reporter plasmid, *Cetn1* gRNA and second half of Cas9 TBD-Cas4.5_N_ under the specific promoter activated during EMT – under the control of TWIST1, resulted in *EGFP* expression. 24h after transfection cells were exposed to TGFβ (10 ng/mL) for 72 h to evaluate the expression level of EMT-related gene *TWIST1*. Next day the monolayer of cells was scratched to form a linear wound. The Wound-healing Assay images were performed up to 72h under Nikon T*i* Eclipse confocal microscope.

### Electrofusion

For cellular fusion was used the BTX’s ECM 380 ElectroCellManipulator. Condition were optimized, two settings of device voltage was used and of controlled electrical pulse: 700V for 15 µs (3 pulses) and 300 V for 500 µs (1 pulse), acting on the cells. For cell electroporation was used hypotonic medium. After experiment cells were resuspended in nutrition medium.

### Confocal microscopy

The cells were cultured on glass bottom plates suitable for confocal imaging. Visualization has been carried out using Nikon T*i* Eclipse microscopy. The nuclei of cells before imaging have been Hoechst 33342 stained (concentration 10 µg/mL).

## RESULTS

Naturally split DnaE intein (DnaE_N_ and DnaE_C_) has high self-affinity followed by efficient splicing and ligation of the flanking exteins ^37^ but only when DnaE_N_ ends with a Cysteine (Cys;C), while DnaE_C_ begins with the following di-amino acids: Cys followed by either Tyr (CY), Trp (CW), Phe (CF), His (CH) or Met (CM)^41^. We then analyzed the Cas9 amino acid sequence and we found only one CY [amino acids (aa) 119-120] and one CF (aa 613-614) (sequences in Supplementary material), since the latter is close to the middle, it was selected, and the two resulting halves have been termed: Cas4.5_N_ (aa 1-613) and Cas4.5_C_ (aa 613-1423). Inteins can spontaneously self-assemble and induce protein ligation of their fused cargoes - in this case, the complementary Cas4.5 fragments, generating the holo-Cas9 endonuclease. To detect Cas9 activity we adopted and modified the EGxxFP reporter^33^. The reporter is constructed on the basis of truncated but overlapping regions of the for enhanced green fluorescent protein (*EGFP*) gene, with an early stop codon after 2/3 of the coding sequence, followed by a genomic (non-coding) fragment from the *Cetn1* gene (stuffer) and a second fragment of the fluorescent protein covering about 500bp of homology and the missing 1/3 of the respective gene. In the presence of active Cas9 and a targeting gRNA to the stuffer region, the homology regions induce homologous recombination, creating the full-length *EGFP* gene and the fluorescent protein being expressed (**Fig. 1**). Co-expression of the 2 halves of Cas9 lacking inteins do not form a holo-enzyme as no reporter activity is detected (not shown).

**Figure 1.**
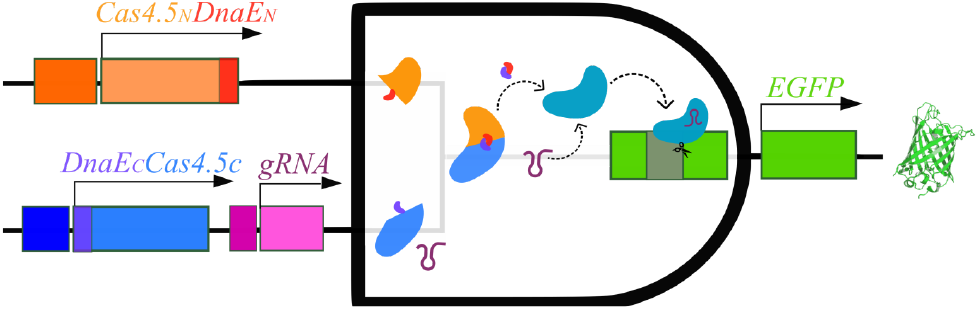
Scheme of the principle of split-Cas9 as an AND logic gate.

Expression of individual Cas4.5 halves in cells, together with the gRNA, showed no reporter activity, proving that they are innocuous on their own. However then both halves, containing inteins, were expressed in the same cell, Cas9 was active as detected by recombination of the reporter.

We then created an AND logic gate where one input is under a conditional promoter, while the other halve is under the ubiquitous (_*p*_*CMV*) promoter (**Fig. 1**). Parts of the system depends on specific promoters and other genetic elements for well-defined subpopulations. Based on that, we are able to program in what kind of cells the system will be activated.

### Cell origin sensing system

As a proof-of-concept, spilt Cas9 was designed to selectively sense cancer cells of epithelial origin, using an epithelial cancer-specific promoter (_*p*_*hCEA*). Carcinoembryonic antigen (*CEA*) has been previously used as a cancer marker due to its selective expression^42,43^, and its promoter has been proposed as cancer-specific and therapeutically useful^44^, thus based on that we cloned *Cas4*.*5*_*N*_*-DnaE*_*N*_ under the _*p*_*hCEA*. Then, the construct was tested on multiple cell lines and based on their known origins we expected expression of *EGFP* only in epithelial cancer cells. Co-transfection of this construct with EGxxFP reporter plasmid, *Cetn1* gRNA and the second half of Cas9 (*DnaE*_*C*_*-Cas4*.*5*_*C*_) driven by a ubiquitous promoter resulted in *EGFP* expression in a pattern matching the cell lines’ origins. Only cancer cell lines of an epithelial origin had all elements expressed and thus *EGFP* expression, which means that all elements have been expressed and once the holo-Cas9 is formed it efficiently activates the reporter. The results confirmed selective expression _*p*_*hCEA*-driven construct **(Fig. 2)**. Our results are in line with the known origins of the studied cell lines. In H1299 human non-small cell lung carcinoma, colon cancer cell line SW480, malignant melanoma cell line A375, and cervical carcinoma cell line HeLa, of epithelial origin we observed expression of reporter (EGFP). In human normal colon (CCD 841 CoTr) cells there was no reporter activity.

**Figure 2.**
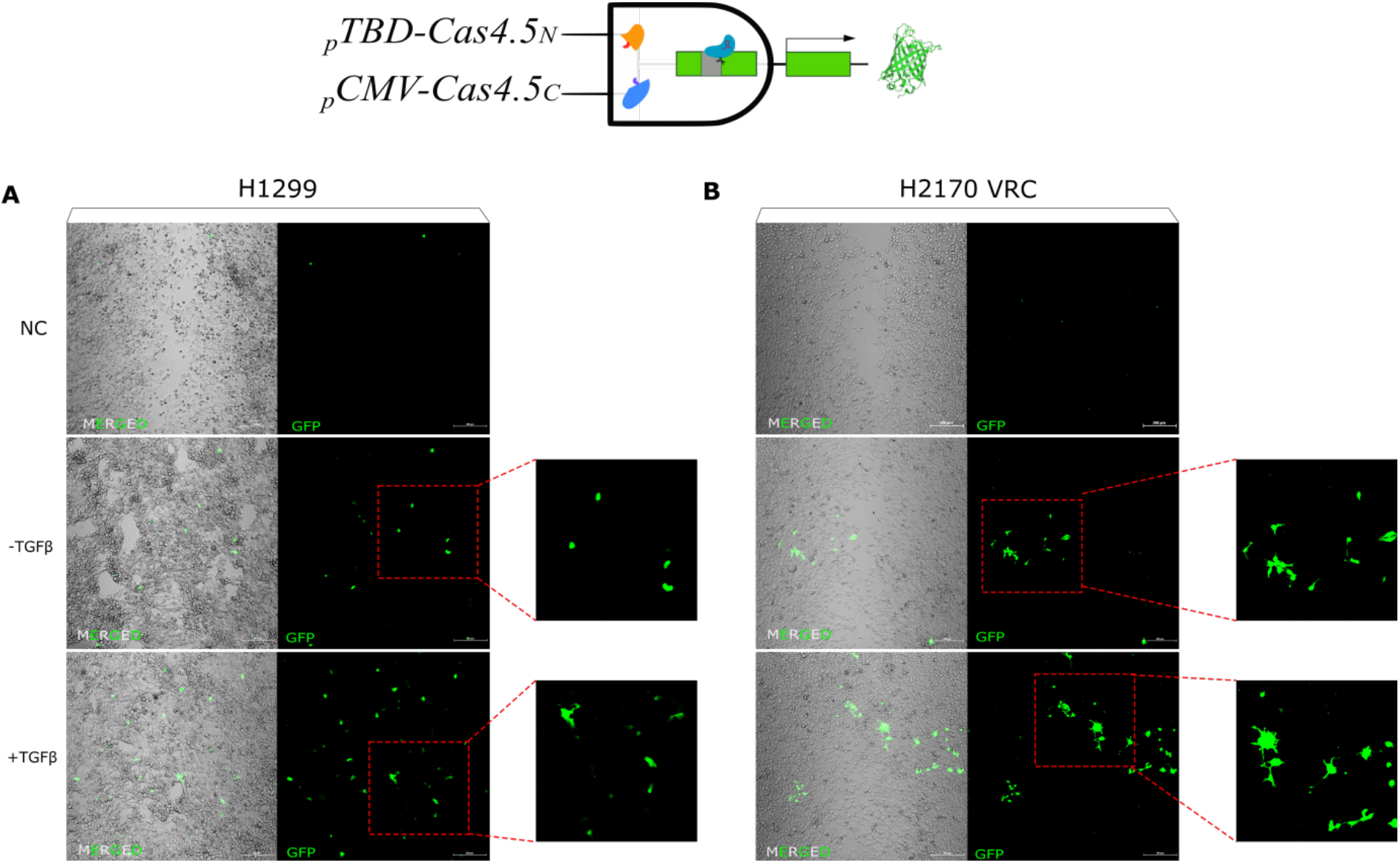
Logic gate to identify only cancer cells of epithelial origin using halves of split Cas9 under a universal (*pCMV*) and a cell type-specific (*phCEA*) promoters. The expression of the green fluorescent protein reporter is observed only in cancer cells of epithelial origin. All cells were co-transfected with _*p*_*hCEA-Cas4*.*5*_*N*_, *pCMV-Cas4*.*5*_*C*_/*gRNA*, the EGxxFP reporter and *pCMV-mCherry* (transfection control) constructs. Co-expression of the split Cas9 halves resulted in holo-enzyme formation via inteins, followed by Cas9-induced homologous recombination of EGFP. Only cancerous cells of epithelial origin expressed both halves since the expression of Cas4.5_N_ is restricted to these cells. Cell lines numbered 1-4 are of epithelial origin and expressed *EGFP*, while control (C, human normal epithelial colon cell line) is not.

Knowing that a cell type, based on its gene expression characteristics, could be labelled, we decided to create another logic gate where a biological process is sensed - in this particular case, epithelial-mesenchymal transition (EMT). For this, we exchanged the universal promoter (_*p*_*CMV*) before *Cas4*.*5*_*C*_*-DnaE*_*C*_ to a promoter that is activated by one of the essential EMT factors, TWIST1^45,46^. TWIST1 activity is closely associated with EMT, and correlates with tumor growth, metastasis and drug resistance, thus diminishing the survival of cancer patients^47^. Using an artificial promoter containing TWIST1 binding domains (*TWIST1-BD*)^35^, and the other halve under the _*p*_*hCEA*, only cells undergoing EMT will express both Cas4.5 halves. To trigger EMT, we exposed cells to TGFβ. Using the wound healing assay, 48h after the addition of TGFβ, we began to observe reporter activity in epithelial carcinoma cells (H1299) and H2170 VRCs cells **(Fig. 3a-b)**, mostly at the edge of the wound as it is often reported^48,49^. The cell morphology, the number of protrusions, on cells that became mesenchymal are clearly visible thanks to the reporter (**Fig. 3b**).

**Figure 2.**
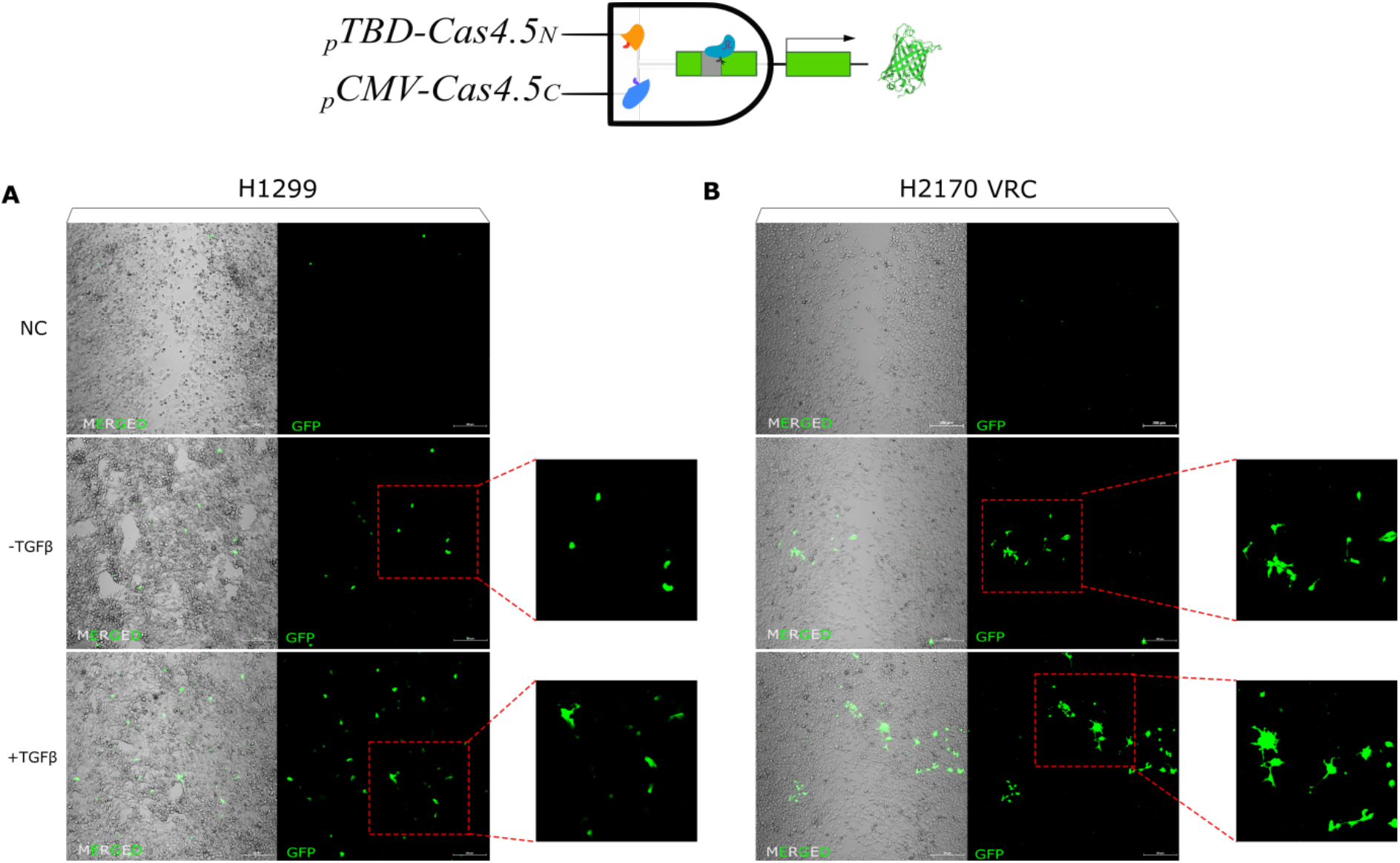
Logic gate for sensing cells that are undergoing EMT. Representative images of H1299 lung cancer cells **(a)** and H2170 VRCs cells **(b)**, both partly mesenchymal. Both cell lines, when exposed to TGFβ as an EMT enhancer, increased the number of *EGFP* expressing cells as a proof of undergoing EMT. Morphological changes were visible after activation of TGFβ showed features of cells of mesenchymal origin - more spindle-shaped, they had more protrusions facilitating migration in comparison to controls.

### Detection of cell-cell fusion events

In addition of changes of gene expression in biological processes, there are other processes where two inputs could be detected by split Cas9 logic gate, such as cell fusion. This is both a physiological, e.g., placentation and skeletal muscle formation, and pathological as in cancer^50,51^, viral HIV^52,53^ or CoV2 infections^54,55^.

To detect cell-cell fusion, two groups of cells, were transfected separately with a different halve of Cas9, plus the gRNA and reporter. The logic gate here is that the inputs, *N*- or *C*-Cas4.5, can only be activated when two different cells fuse to each other, sharing a common cytoplasm.

To induce fusion, we applied two methods: the overexpression of a “fusogene” Syncytin1 - a protein involved in syncytium formation in placenta^56^ that is known to promote the formation of multinucleated cells (syncytia)^57,58^; or a chemical-based method where polyethylene glycol (PEG), due to its known fusogenic properties^59^.

For this purpose we used transiently transfected cell lines (HEK293, H1299, H2170), each line expressing only one of the halves of Cas9 at a time (Cas4.5_C_ or Cas4.5_N_), and co-transfected with the reporter (EGxxFP), as control cells, or either co-transfected with Syncytin1 plasmid or PEG-treated to induce cell-cell fusion. 24 hours after transfection, cells expressing one half were admixed with either the same lineage, but expressing the complementing Cas4.5, or with a different cell type but expressing the complementing Cas4.5. As expected, in control samples cell fusion was undetectable, while a number of fusion events occurred, in particular in mixed cells HEK293+H1299 and HEK293+H2170, in those either expressing Syncytin1 or PEG-treated (**Fig. 4a-b**). However, treatment with PEG showed toxic effects on cells and therefore we continued only with Syncytin1 overexpression for further experiments. We then tested electrofusion – the use of high voltage electrical pulses to induce fusions of cells in close proximity^60^ – on an admixture of the cells expressing either halve. For this aim, different voltages were tested: 300V 500 µs one pulse vs 700V 15 µs three pulses. Compared to control samples, where cell fusion was undetectable (**Fig.4 c,f**), in cells expressing both halves, electrofusion resulted in a few fusion events (**Fig. 4d,g**). Thus, to enhance fusion, we then combined both approaches, electrofusion and either expression of Syncytin1 (**Fig. 4e,h**). As a result, 700V stimulation together with Syncytin1 induced fusion more effectively than the 300V, in combination of different cell types HEK293 + H1299 (**Fig. 4e**), and HEK293 + H2170 (**Fig. 4g**). We show that in a mix of cells HEK293+H2170, Syncytin1 overexpression alone and in pair with electrofusion visibly increased fusion events (**Fig. 4b,h**).

**Figure 4.**
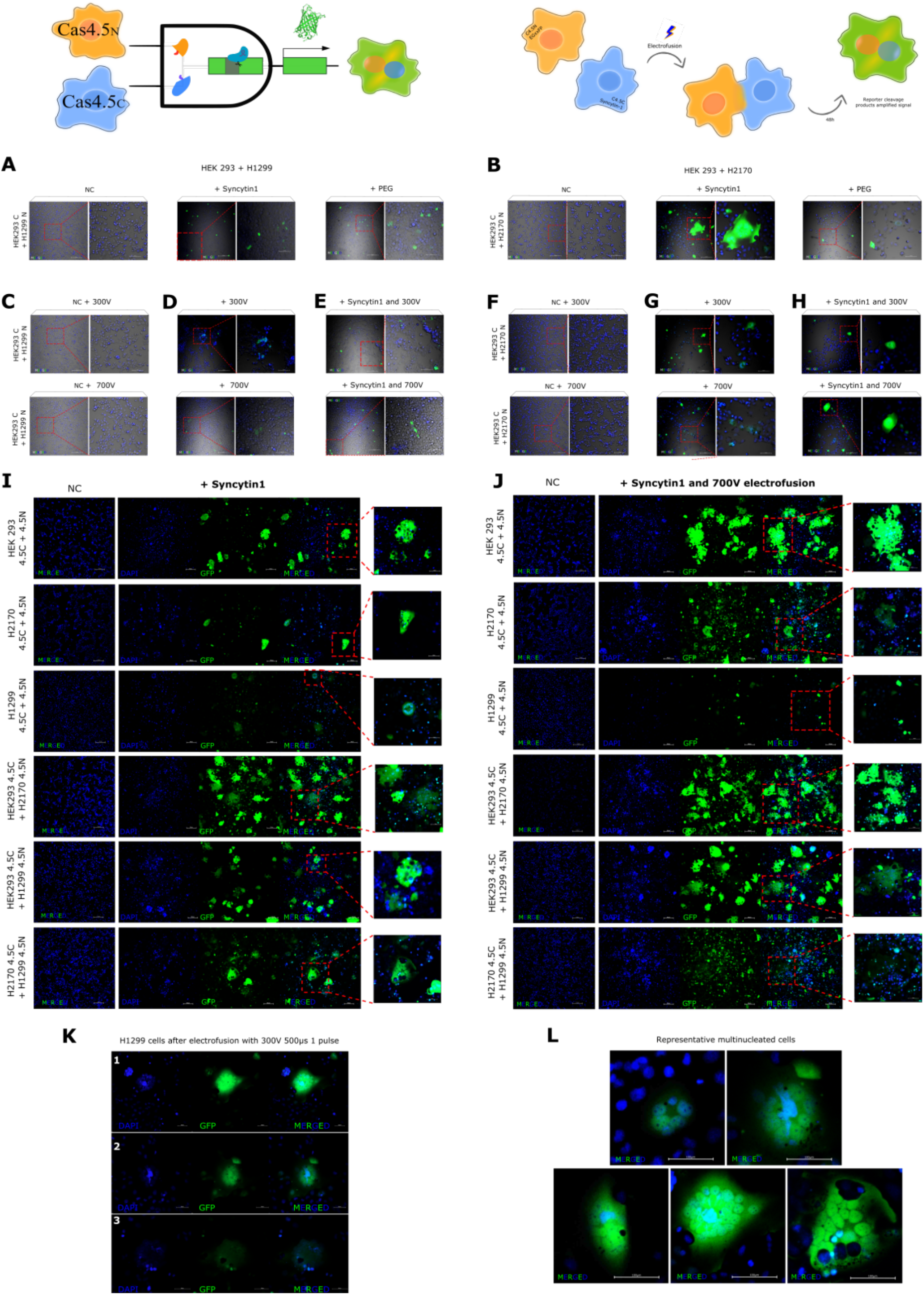
Logic gate to detect cell-cell fusion and schematic diagram of cell-cell electrofusion and hybrid cell formation. Images of admixed transiently transfected cells with complementing Cas9 halves HEK 293 + H1299 and HEK 293 + H2170 after fusion induction by overexpression of Syncytin-1, alone or in pair with electrofusion, or by PEG treatment. Control cells (NC) were transfected with only one half of Cas9 plus the reporter and fusogenic vectors (**a-b**). As we expected admixed cells HEK293 + H1299 expressing either one halve (as a control samples) and electrofused using 300V and 700V do not show any fusion events (**c**), compared to electrofusion of cells expressing both halves where we got results of a few cell fusion (**d**). After enhancement of fusion by electric pulses in pair with Syncytin1 overexpression, we visibly increased occurrence of fusion events (**e**). In control samples of admixed HEK293 + H2170 cells after electrofusion using 300 V and 700 V, also the fusion was undetectable (**f**), in comparison to Syncytin1-induced fusion cells plus electrofusion, where fusion cell were clearly visible (**h**). Images of admixed stable cell lines with complementing Cas9 halves (HEK293, H1299, H2170, HEK293+H1299, HEK293+H2170, and H1299+H2170) after fusion induction by overexpression of Syncytin1, and Syncytin1 plus electrofusion. Control cells (NC) were transfected with only one half of Cas9 plus the reporter and fusogenic vectors. In relation to cells only expressed fusogenic Syncytin1 (**i**), the combination of Syncytin1 and electrical pulses (**j**) greatly enhanced the detection of syncytia. When fusing HEK293 with H1299 or H2170, the syncytia visibly formed contained more nuclei, than lung cancer cells in a mix within the same line. H1299 cells overexpressing Syncytin1 in pair with electrofusion at 700V did not show significant fusion events (**i-j**). Higher efficiency was achieved by expression of Syncytin1 and lower voltage (300 V 500 µs one pulse) in stable expressing complementing Cas9 halves H1299 cells **(k)**. Representative large multinucleated syncytia after electrofusion cells overexpressed Syncytin1 H1299 (using 300 V), H2170 cell lines (using 700V), and mixed HEK 293 with H1299 cell line and HEK293 with H2170 cells (using 700V) **(l)**.

**Figure 5.**
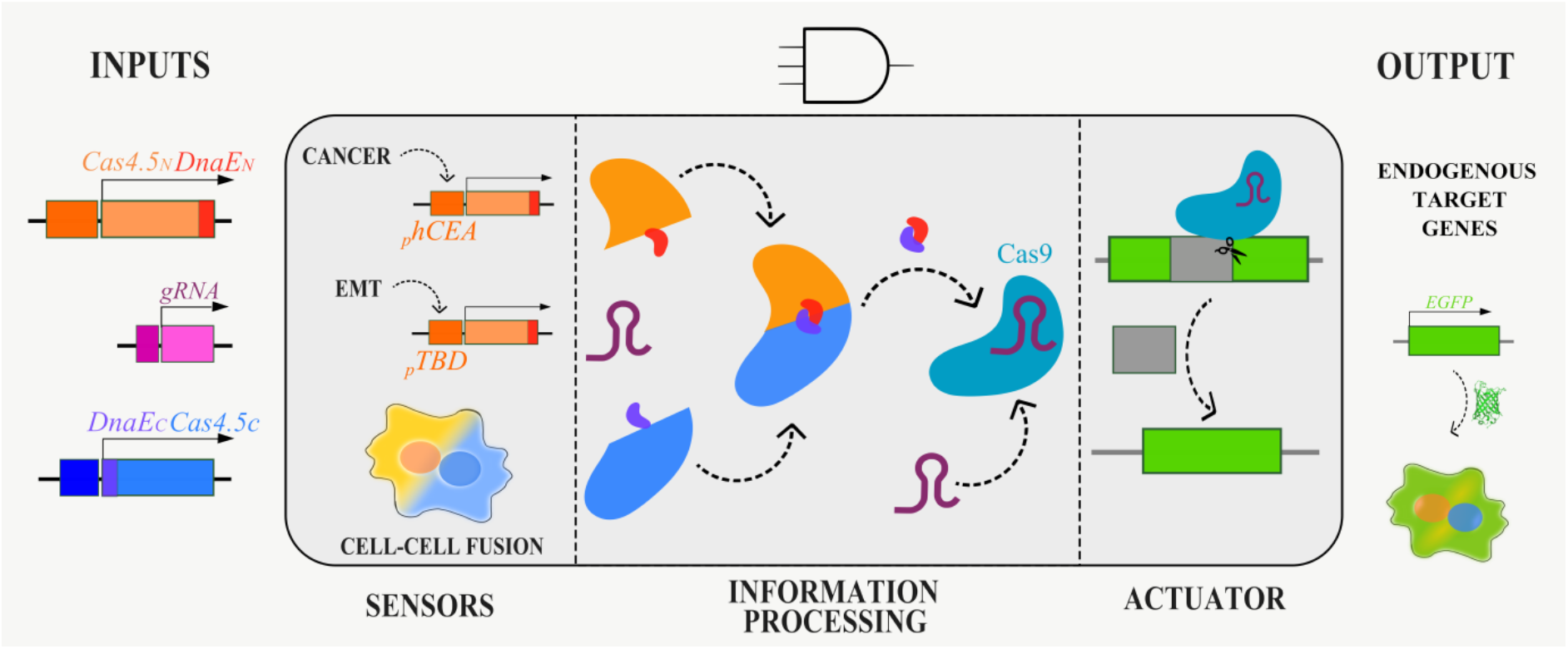
Engineering cells with CRISPR/Cas9 circuits.

For more in-depth inquiry, we created stable cell lines, and used optimized settings to induce fusion of different cell, each line stably expressing one half of Cas9 (Cas4.5_C_ or Cas4.5_N_), and co-transfected with the reporter (EGxxFP) and Syncytin1 plasmids. 24 hours after transfection, the different cell types were admixed within either the same lineage, but expressing the complementing Cas4.5, or to a different cell type also expressing the complementing Cas4.5 (HEK293 with H2170, HEK293 with H1299, and H2170 with H1299) and then cells were electrofused (700 V for 15 µs). Control cells were co-transfected with only one half of Cas9 plus the reporter and Syncytin1 plasmids (negative control, NC). Compared to cells only overexpressing Syncytin1 (**Fig. 4i**), the presence of Syncytin1 and electrical pulses (**Fig. 4j**) greatly enhanced the detection of syncytia. We show that in a mix of different type of cells (when fusing HEK293 with H1299 or H2170) the syncytia formed were larger and contained more nuclei, than either cell type fused with its same lineage (**Fig. 4i-j**). There were no major fusion events in the mix of H1299 cells, whether only Syncytin1 was overexpressed (**Fig. 4i**) or in pair with 700 V stimulation (**Fig. 4j**). Cells survival was lower after 1 long vs 3 shorter pulses. Surprisingly, the H1299 cells generated more fusion events at lower voltage (300 V for 500 µs) and 1 pulse (**Fig. 4k**). The presence of Syncytin1 and electrical impulses contributed significantly for syncytia to occur in H1299 (using 300 V) and H2170 cell lines (using 700V), and mixed HEK 293 with H1299 or HEK293 with H2170 cells (using 700V) **(Fig. 4l)**.

## DISCUSSION

Prokaryotes, which include bacteria and archaea, have simpler signaling processes than eukaryotic cells, which means that SynBio logic gates are easier to generate. In mammalian systems signaling is rather complex and therefore simple SynBio circuits are more demanding to design.

While biological processes or events can be detected using biochemical methods, the use of bioassays have become more and more commonplace. Yet, many bioassays contain single inputs and can hardly be adapted to other purposes. Here, we show that it is possible to generate logic gates using a split CRISPR/Cas9, which is an actuator with many downstream options^61–64^. Moreover, the activation of Cas9 also creates a memory so using this system we could detect biological processes after they occurred e.g. *in vivo* long after they occurred.

As CRISPR-Cas-mediated genome editing technologies have provided accessible and flexible means to alter, regulate and visualize genomes, they are considered a milestone for molecular biology in the 21st century. So far, CRISPR-Cas systems have found wide applications in the investigation of gene function, gene therapy, development of targeted drugs, and construction of animal models, which fully proves their great potential for further development^10,65–68^.

Together with other synthetic systems for detecting biological phenomena such as optogenetic, gene expression and cell-free sensors, CRISPR systems have shown incredible flexibility that can be easily translated to various research and medical contexts. For example, Liu *et al*. reported a modular AND logic gate based on the CRISPR-Cas9 system, contributing to a synthetic biology platform for targeting and controlling bladder cancer cells *in vitro*^27^.

Here, we introduced a new application of the CRISPR/Cas9, as a sensor for detection cancer cells with epithelial origin, using an epithelial cancer-specific promoter (_*p*_*hCEA*), a biological process - cells undergoing EMT, The selection of the *phCEA*, a marker to detect the presence of cancer cells of epithelial origin is specific for various types of cancer including colorectal cancer^44,69^, supports the concept that CEA is one of the most promising target antigens for colorectal cancer immunotherapy in use. CEA-targeted immunotherapies are presently in the preclinical or clinical phase, including CEA vaccines and CEA-targeted CAR T cells or TCR-modified T cells^70^.

Finally, split Cas9 was used for the detection of cell-cell fusions. A proces that has been reported to occur between tumor cells and surrounding cells^71–73^. Where normal cells surrounding cancer cells can acquire malignant characteristics, according to the proverb "lie down with dogs, rise up with fleas”. One of the mechanisms facilitating tumor progression as the tumor spreads to surrounding tissue is cell-cell fusion^74^. Cell-cell fusion has been suggested as one possible mechanism for tumor progression^75,76^. Given that EMT is closely related to tumor invasiveness, there is likely to be a link between cell fusion and tumor invasiveness. When cancer cells fuse with healthy differentiated cells, the resulting hybrid cell can develop the ability to divide uncontrollably and form tumours^77–79^. The most compelling evidence that cell-cell fusion drives cancerogenesis is the report by Lazova and collaborators^80^ where they found that a metastatic melanoma was a hybrid between the patient’s cells and the ones from his brother after an allogeneic bone-marrow transplantation. Since then, more reports of cell-cell fusion show that such events initiate or fuel tumorigenesis^81,82^.

The applications presented above are additional tools to the CRISPR palette. By modifying the genetic elements that control the expression of its individual elements, it can be programmed and created more complex logic gates. One of the possibilities offered by this system is the elimination of a selected cell subpopulation from a given cell mixture through the use of gRNAs directed against essential genes. Thus, the proposed method may have potential for cancer diagnosis, therapy and treatment monitoring.

In recent years, the potential of using Cas9 as an actuator has also become apparent, is still a relatively new approach, but it has the potential to enable new possibilities for precise control over biological systems. Moreover, the requirement of the gRNA allows a 3-input system that can target specific genomic loci or additional logic gates/circuits, and thus become a research tool for multiple applications.

In the last few years, SynBio has made evident that it can use engineering logic to provide insight into previously unrecognized mechanisms of natural phenomena^83,84^. As Cas9 has recently been shown to be associated with non-canonical RNAs in bacteria^83^, exploring the interplay among endogenous RNA and Cas9 in eukaryotes could point to new technology, including additional imaging reporters, to study gene activation and even enables therapy modalities in mammalian systems.

In the grand scheme, CRISPR provides new opportunities for constructing circuits that control processes in living cells with wide application in synthetic biology, but there are still limitations that need to be overcome. Our findings have expanded the use of CRISPR technologies to more complex gates in the future.

## AUTHOR CONTRIBUTIONS

AP-P performed the experiments and analyses. JCz assisted with specific promotors experiments and data analyses. JK assisted in the fusion experiment and helped writing the manuscript. AP-P, JCz, and AR-M designed the experimental section of this research work and co-wrote the manuscript draft. AR-M supervised the overall study. All authors read and approved the final manuscript.

## ACKNOWLEDGEMENTS

Authors would like to thank Addgene, and the colleagues depositing their plasmids there, for the support to the greater scientific community.

