## Supplementary material for "Synthetic Circuits Based on Split CAS9 to Detect Cellular Events": Suppl. Material

Phone No.: +48 (81) 4486363

**Statements**:

All the raw data supporting the findings in this work can be obtained on request from the corresponding author.

Plasmids used in this work will be available via Addgene and on request from the corresponding author.

**Nucleic acid Sequence of the construct $_{p}$CMV-Cas4.5$_{N}$-DnaE$_{N}$ (7054 bp)**

gagggcctatttcccatgattccttcatatttgcatatacgatacaaggctgttagagagataattggaattaatttgactgtaaacacaaag
atattagtacaaaatacgtgacgtagaaagtaataatttcttgggtagtttgcagttttaaaattatgttttaaaatggactatcatatgcttacc
gtaacttgaaagtatttcgatttcttggctttatatatcttGTGGAAAGGACGAAACACCggGTCTTCgaGAAG
ACCTGTTTTAGAGCTAGAAATAGCAAGTTAAAATAAGGCTAGTCCGTTATCAACT
TGAAAAAGTGGCACCGAGTCGGTGcTTTTTTgttttagagctagaaatagcaagttaaaataaggctagtc
cgtTTTTagcgcgtgcgccaattctgcagacaaatggctctagaggtacccgttacataacttacggtaaatggcccgcctggctga
ccgcccaacgacccccgcccattgacgtcaatagtaacgccaatagggactttccattgacgtcaatgggtggagtatttacggtaaac
tgcccacttggcagtacatcaagtgtatcatatgccaagtacgccccctattgacgtcaatgacggtaaatggcccgcctggcattGtg
cccagtacatgaccttatgggactttcctacttggcagtacatctacgtattagtcatcgctattaccatggtcgaggtgagccccacgttc
tgcttcactctccccatctcccccccctccccacccccaattttgtatttatttattttttaattattttgtgcagcgatgggggcgggggggg
ggggggggcgcgcgccaggcggggcggggcggggcgaggggcggggcggggcgaggcggagaggtgcggcggcagcca
atcagagcggcgcgctccgaaagtttccttttatggcgaggcggcggcggcggcggccctataaaaagcgaagcgcgcggcggg
cgggagtcgctgcgacgctgccttcgccccgtgccccgctccgccgccgcctcgcgccgcccgccccggctctgactgaccgcgtt
actcccacaggtgagcgggcgggacggcccttctcctccgggctgtaattagctgagcaagaggtaagggtttaagggatggttggtt
ggtggggtattaatgtttaattacctggagcacctgcctgaaatcactttttttcaggttGGaccggtgccaccATGGACTATA
AGGACCACGACGGAGACTACAAGGATCATGATATTGATTACAAAGACGATGACG
ATAAGATGGCCCCAAAGAAGAAGCGGAAGGTCGGTATCCACGGAGTCCCAGCA
GCCGACAAGAAGTACAGCATCGGCCTGGACATCGGCACCAACTCTGTGGGCTGG
GCCGTGATCACCGACGAGTACAAGGTGCCCAGCAAGAAATTCAAGGTGCTGGGC
AACACCGACCGGCACAGCATCAAGAAGAACCTGATCGGAGCCCTGCTGTTCGAC
AGCGGCGAAACAGCCGAGGCCACCCGGCTGAAGAGAACCGCCAGAAGAAGATA
CACCAGACGGAAGAACCGGATCTGCTATCTGCAAGAGATCTTCAGCAACGAGAT
GGCCAAGGTGGACGACAGCTTCTTCCACAGACTGGAAGAGTCCTTCCTGGTGGA
AGAGGATAAGAAGCACGAGCGGCACCCCATCTTCGGCAACATCGTGGACGAGGT
GGCCTACCACGAGAAGTACCCCACCATCTACCACCTGAGAAAGAAACTGGTGGA
CAGCACCGACAAGGCCGACCTGCGGCTGATCTATCTGGCCCTGGCCCACATGATC
AAGTTCCGGGGCCACTTCCTGATCGAGGGCGACCTGAACCCCGACAACAGCGAC
GTGGACAAGCTGTTCATCCAGCTGGTGCAGACCTACAACCAGCTGTTCGAGGAA
AACCCCATCAACGCCAGCGGCGTGGACGCCAAGGCCATCCTGTCTGCCAGACTG
AGCAAGAGCAGACGGCTGGAAAATCTGATCGCCCAGCTGCCCGGCGAGAAGAA
GAATGGCCTGTTCGGAAACCTGATTGCCCTGAGCCTGGGCCTGACCCCCAACTTC
AAGAGCAACTTCGACCTGGCCGAGGATGCCAAACTGCAGCTGAGCAAGGACACC
TACGACGACGACCTGGACAACCTGCTGGCCCAGATCGGCGACCAGTACGCCGAC
CTGTTTCTGGCCGCCAAGAACCTGTCCGACGCCATCCTGCTGAGCGACATCCTGA
GAGTGAACACCGAGATCACCAAGGCCCCCCTGAGCGCCTCTATGATCAAGAGAT
ACGACGAGCACCACCAGGACCTGACCCTGCTGAAAGCTCTCGTGCGGCAGCAGC
TGCCTGAGAAGTACAAAGAGATTTTCTTCGACCAGAGCAAGAACGGCTACGCCG
GCTACATTGACGGCGGAGCCAGCCAGGAAGAGTTCTACAAGTTCATCAAGCCCA
TCCTGGAAAAGATGGACGGCACCGAGGAACTGCTCGTGAAGCTGAACAGAGAG
GACCTGCTGCGGAAGCAGCGGACCTTCGACAACGGCAGCATCCCCCACCAGATC
CACCTGGGAGAGCTGCACGCCATTCTGCGGCGGCAGGAAGATTTTTACCCATTCC
TGAAGGACAACCGGGAAAAGATCGAGAAGATCCTGACCTTCCGCATCCCCTACT
ACGTGGGCCCTCTGGCCAGGGGAAACAGCAGATTCGCCTGGATGACCAGAAAGA
GCGAGGAAACCATCACCCCCTGGAACTTCGAGGAAGTGGTGGACAAGGGCGCTT
CCGCCCAGAGCTTCATCGAGCGGATGACCAACTTCGATAAGAACCTGCCCAACG
AGAAGGTGCTGCCCAAGCACAGCCTGCTGTACGAGTACTTCACCGTGTATAACG
AGCTGACCAAAGTGAAATACGTGACCGAGGGAATGAGAAAGCCCGCCTTCCTGA
GCGGCGAGCAGAAAAAGGCCATCGTGGACCTGCTGTTCAAGACCAACCGGAAAG

TGACCGTGAAGCAGCTGAAAGAGGACTACTTCAAGAAAATCGAGGGATCCTGTT
TAAGCTATGAAACGGAAATATTGACAGTAGAATATGGATTATTACCGATTGGTA
AAATTGTAGAAAAGCGCATCGAATGTACTGTTTATAGCGTTGATAATAATGGAA
ATATTTATACACAACCTGTAGCACAATGGCACGATCGCGGAGAACAAGAGGTGT
TTGAGTATTGTTTGGAAGATGGTTCATTGATTCGGGCAACAAAAGACCATAAGTT
TATGACTGTTGATGGTCAAATGTTGCCAATTGATGAAATATTTGAACGTGAATTG
GATTTGATGCGGGTTGATAATTTGCCGAATgaattcGGCAGTGGAGAGGGCAGAGG
AAGTCTGCTAACATGCGGTGACGTCGAGGAGAATCCTGGCCCAatgaccgagtacaagccc
acggtgcgcctcgccacccgcgacgacgtccccagggccgtacgcaccctcgccgccgcgttcgccgactaccccgccacgcgc
cacaccgtcgatccggaccgccacatcgagcgggtcaccgagctgcaagaactcttcctcacgcgcgtcgggctcgacatcggcaa
ggtgtgggtcgcggacgacggcgccgcggtggcggtctggaccacgccggagagcgtcgaagcgggggcggtgttcgccgaga
tcggcccgcgcatggccgagttgagcggttcccggctggccgcgcagcaacagatggaaggcctcctggcgccgcaccggccca
aggagcccgcgtggttcctggccaccgtcggcgtctcgcccgaccaccagggcaagggtctgggcagcgccgtcgtgctccccgg
agtggaggcggccgagcgcgccggggtgcccgccttcctggagacctccgcgccccacaacctccccttctacgagcggctcggc
ttcaccgtcaccgccgacgtcgaggtgcccgaaggaccgcgcacctggtgcatgacccgcaagcccggtgcctgagaattctaaC
TAGAGCTCGCTGATCAGCCTCGACTGTGCCTTCTAGTTGCCAGCCATCTGTTGTTT
GCCCCTCCCCCGTGCCTTCCTTGACCCTGGAAGGTGCCACTCCCACTGTCCTTTCC
TAATAAAATGAGGAAATTGCATCGCATTGTCTGAGTAGGTGTCATTCTATTCTGG
GGGGTGGGGTGGGGCAGGACAGCAAGGGGGAGGATTGGGAAGAgAATAGCAGG
CATGCTGGGGAgcggccgcaggaacccctagtgatggagttggccactccctctctgcgcgctcgctcgctcactgaggc
cgggcgaccaaaggtcgcccgacgcccgggctttgcccgggcggcctcagtgagcgagcgagcgcgcagctgcctgcaggggc
gcctgatgcggtattttctccttacgcatctgtgcggtatttcacaccgcatacgtcaaagcaaccatagtacgcgccctgtagcggcgc
attaagcgcggcgggtgtggtggttacgcgcagcgtgaccgctacacttgccagcgccctagcgcccgctcctttcgctttcttcccttc
ctttctcgccacgttcgccggctttccccgtcaagctctaaatcgggggctccctttagggttccgatttagtgctttacggcacctcgacc
ccaaaaaacttgatttgggtgatggttcacgtagtgggccatcgccctgatagacggtttttcgccctttgacgttggagtccacgttcttt
aatagtggactcttgttccaaactggaacaacactcaaccctatctcgggctattcttttgatttataagggattttgccgatttcggcctatt
ggttaaaaaatgagctgatttaacaaaaatttaacgcgaattttaacaaaatattaacgtttacaattttatggtgcactctcagtacaatctg
ctctgatgccgcatagttaagccagccccgacacccgccaacacccgctgacgcgccctgacgggcttgtctgctcccggcatccgc
ttacagacaagctgtgaccgtctccgggagctgcatgtgtcagaggttttcaccgtcatcaccgaaacgcgcgagacgaaagggcct
cgtgatacgcctatttttataggttaatgtcatgataataatggtttcttagacgtcaggtggcacttttcggggaaatgtgcgcggaaccc
ctatttgtttatttttctaaatacattcaaatatgtatccgctcatgagacaataaccctgataaatgcttcaataatattgaaaaaggaagagt
atgagtattcaacatttccgtgtcgcccttattcccttttttgcggcattttgccttcctgtttttgctcacccagaaacgctggtgaaagtaaa
agatgctgaagatcagttgggtgcacgagtgggttacatcgaactggatctcaacagcggtaagatccttgagagttttcgccccgaag
aacgttttccaatgatgagcacttttaaagttctgctatgtggcgcggtattatcccgtattgacgccgggcaagagcaactcggtcgcc
gcatacactattctcagaatgacttggttgagtactcaccagtcacagaaaagcatcttacggatggcatgacagtaagagaattatgca
gtgctgccataaccatgagtgataacactgcggccaacttacttctgacaacgatcggaggaccgaaggagctaaccgcttttttgcac
aacatgggggatcatgtaactcgccttgatcgttgggaaccggagctgaatgaagccataccaaacgacgagcgtgacaccacgatg
cctgtagcaatggcaacaacgttgcgcaaactattaactggcgaactacttactctagcttcccggcaacaattaatagactggatggag
gcggataaagttgcaggaccacttctgcgctcggcccttccggctggctggtttattgctgataaatctggagccggtgagcgtggaag
ccgcggtatcattgcagcactggggccagatggtaagccctcccgtatcgtagttatctacacgacggggagtcaggcaactatggat
gaacgaaatagacagatcgctgagataggtgcctcactgattaagcattggtaactgtcagaccaagtttactcatatatactttagattg
atttaaaacttcatttttaatttaaaaggatctaggtgaagatcctttttgataatctcatgaccaaaatcccttaacgtgagttttcgttccact
gagcgtcagaccccgtagaaaagatcaaaggatcttcttgagatcctttttttctgcgcgtaatctgctgcttgcaaacaaaaaaaccacc
gctaccagcggtggtttgtttgccggatcaagagctaccaactctttttccgaaggtaactggcttcagcagagcgcagataccaaatac
tgtccttctagtgtagccgtagttaggccaccacttcaagaactctgtagcaccgcctacatacctcgctctgctaatcctgttaccagtgg
ctgctgccagtggcgataagtcgtgtcttaccgggttggactcaagacgatagttaccggataaggcgcagcggtcgggctgaacgg
ggggttcgtgcacacagcccagcttggagcgaacgacctacaccgaactgagatacctacagcgtgagctatgagaaagcgccacg
cttcccgaagggagaaaggcggacaggtatccggtaagcggcagggtcggaacaggagagcgcacgagggagcttccaggggg
aaacgcctggtatctttatagtcctgtcgggtttcgccacctctgacttgagcgtcgatttttgtgatgctcgtcaggggggcggagcctat
ggaaaaacgccagcaacgcggcctttttacggttcctggccttttgctggccttttgctcacatgt

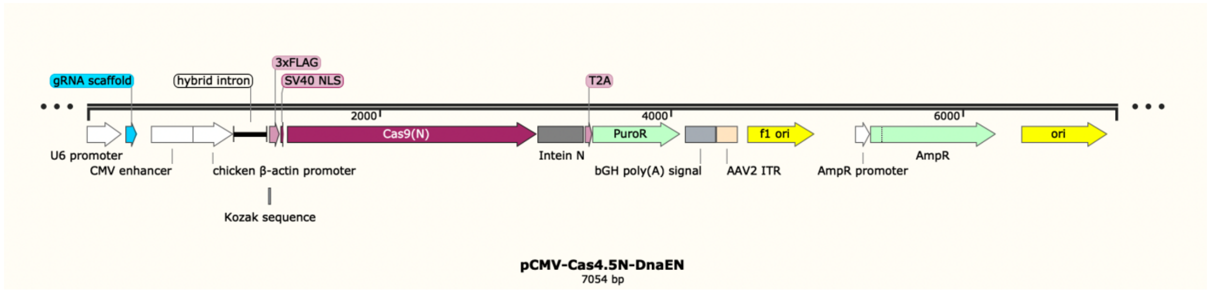


### Nucleic acid Sequence of the construct $_{p}$CMV-Cas4.5$_{C}$-DnaE$_{C}$ (7591 bp)

gagggcctatttcccatgattccttcatatttgcatatacgatacaaggctgttagagagataattggaattaatttgactgtaaacacaaag
atattagtacaaaatacgtgacgtagaaagtaataatttcttgggtagtttgcagttttaaaattatgttttaaaatggactatcatatgcttacc
gtaacttgaaagtatttcgatttcttggctttatatatcttGTGGAAAGGACGAAACACCggGTCTTCgaGAAG
ACCTGTTTTAGAGCTAGAAATAGCAAGTTAAAATAAGGCTAGTCCGTTATCAACT
TGAAAAAGTGGCACCGAGTCGGTGcTTTTTTgttttagagctagaaatagcaagttaaaataaggctagtc
cgtTTTTagcgcgtgcgccaattctgcagacaaatggctctagaggtacccgttacataacttacggtaaatggcccgcctggctga
ccgcccaacgacccccgcccattgacgtcaatagtaacgccaatagggactttccattgacgtcaatgggtggagtatttacggtaaac
tgcccacttggcagtacatcaagtgtatcatatgccaagtacgccccctattgacgtcaatgacggtaaatggcccgcctggcattGtg
cccagtacatgaccttatgggactttcctacttggcagtacatctacgtattagtcatcgctattaccatggtcgaggtgagccccacgttc
tgcttcactctccccatctcccccccctccccacccccaattttgtatttatttattttttaattattttgtgcagcgatgggggcgggggggg
ggggggggcgcgcgccaggcggggcggggcggggcgaggggcggggcggggcgaggcggagaggtgcggcggcagcca
atcagagcggcgcgctccgaaagtttccttttatggcgaggcggcggcggcggcggccctataaaaagcgaagcgcgcggcggg
cgggagtcgctgcgacgctgccttcgccccgtgccccgctccgccgccgcctcgcgccgcccgccccggctctgactgaccgcgtt
actcccacaggtgagcgggcgggacggcccttctcctccgggctgtaattagctgagcaagaggtaagggtttaagggatggttggtt
ggtggggtattaatgtttaattacctggagcacctgcctgaaatcactttttttcaggttGGaccggtgccaccATGGACTATA
AGGACCACGACGGAGACTACAAGGATCATGATATTGATTACAAAGACGATGACG
ATAAGATGGCCCCAAAGAAGAAGCGGAAGGTCGGTcatggtggcggtggctctggaggtggtgg
gtccggaggaggcggccgcatcaaaatagccacacgtaaatatttaggcaaacaaaatgtctatgacattggagttgagcgcgaccat
aattttgcactcaaaaatggcttcatagcttctaatTGCTTCGACTCCGTGGAAATCTCCGGCGTGGAAG
ATCGGTTCAACGCCTCCCTGGGCACATACCACGATCTGCTGAAAATTATCAAGGA
CAAGGACTTCCTGGACAATGAGGAAAACGAGGACATTCTGGAAGATATCGTGCT
GACCCTGACACTGTTTGAGGACAGAGAGATGATCGAGGAACGGCTGAAAACCTA
TGCCCACCTGTTCGACGACAAAGTGATGAAGCAGCTGAAGCGGCGGAGATACAC
CGGCTGGGGCAGGCTGAGCCGGAAGCTGATCAACGGCATCCGGGACAAGCAGTC
CGGCAAGACAATCCTGGATTTCCTGAAGTCCGACGGCTTCGCCAACAGAAACTTC
ATGCAGCTGATCCACGACGACAGCCTGACCTTTAAAGAGGACATCCAGAAAGCC
CAGGTGTCCGGCCAGGGCGATAGCCTGCACGAGCACATTGCCAATCTGGCCGGC
AGCCCCGCCATTAAGAAGGGCATCCTGCAGACAGTGAAGGTGGTGGACGAGCTC
GTGAAAGTGATGGGCCGGCACAAGCCCGAGAACATCGTGATCGAAATGGCCAGA
GAGAACCAGACCACCCAGAAGGGACAGAAGAACAGCCGCGAGAGAATGAAGCG
GATCGAAGAGGGCATCAAAGAGCTGGGCAGCCAGATCCTGAAAGAACACCCCGT
GGAAAACACCCAGCTGCAGAACGAGAAGCTGTACCTGTACTACCTGCAGAATGG
GCGGGATATGTACGTGGACCAGGAACTGGACATCAACCGGCTGTCCGACTACGA
TGTGGACCATATCGTGCCTCAGAGCTTTCTGAAGGACGACTCCATCGACAACAAG
GTGCTGACCAGAAGCGACAAGAACCGGGGCAAGAGCGACAACGTGCCCTCCGA
AGAGGTCGTGAAGAAGATGAAGAACTACTGGCGGCAGCTGCTGAACGCCAAGCT

GATTACCCAGAGAAAGTTCGACAATCTGACCAAGGCCGAGAGAGGCGGCCTGAG
CGAACTGGATAAGGCCGGCTTCATCAAGAGACAGCTGGTGGAAACCCGGCAGAT
CACAAAGCACGTGGCACAGATCCTGGACTCCCGGATGAACACTAAGTACGACGA
GAATGACAAGCTGATCCGGGAAGTGAAAGTGATCACCCTGAAGTCCAAGCTGGT
GTCCGATTTCCGGAAGGATTTCCAGTTTTACAAAGTGCGCGAGATCAACAACTAC
CACCACGCCCACGACGCCTACCTGAACGCCGTCGTGGGAACCGCCCTGATCAAA
AAGTACCCTAAGCTGGAAAGCGAGTTCGTGTACGGCGACTACAAGGTGTACGAC
GTGCGGAAGATGATCGCCAAGAGCGAGCAGGAAATCGGCAAGGCTACCGCCAA
GTACTTCTTCTACAGCAACATCATGAACTTTTTCAAGACCGAGATTACCCTGGCC
AACGGCGAGATCCGGAAGCGGCCTCTGATCGAGACAAACGGCGAAACCGGGGA
GATCGTGTGGGATAAGGGCCGGGATTTTGCCACCGTGCGGAAAGTGCTGAGCAT
GCCCCAAGTGAATATCGTGAAAAAGACCGAGGTGCAGACAGGCGGCTTCAGCAA
AGAGTCTATCCTGCCCAAGAGGAACAGCGATAAGCTGATCGCCAGAAAGAAGGA
CTGGGACCCTAAGAAGTACGGCGGCTTCGACAGCCCCACCGTGGCCTATTCTGTG
CTGGTGGTGGCCAAAGTGGAAAAGGGCAAGTCCAAGAAACTGAAGAGTGTGAA
AGAGCTGCTGGGGATCACCATCATGGAAAGAAGCAGCTTCGAGAAGAATCCCAT
CGACTTTCTGGAAGCCAAGGGCTACAAAGAAGTGAAAAAGGACCTGATCATCAA
GCTGCCTAAGTACTCCCTGTTCGAGCTGGAAAACGGCCGGAAGAGAATGCTGGC
CTCTGCCGGCGAACTGCAGAAGGGAAACGAACTGGCCCTGCCCTCCAAATATGT
GAACTTCCTGTACCTGGCCAGCCACTATGAGAAGCTGAAGGGCTCCCCCGAGGA
TAATGAGCAGAAACAGCTGTTTGTGGAACAGCACAAGCACTACCTGGACGAGAT
CATCGAGCAGATCAGCGAGTTCTCCAAGAGAGTGATCCTGGCCGACGCTAATCT
GGACAAAGTGCTGTCCGCCTACAACAAGCACCGGGATAAGCCCATCAGAGAGCA
GGCCGAGAATATCATCCACCTGTTTACCCTGACCAATCTGGGAGCCCCTGCCGCC
TTCAAGTACTTTGACACCACCATCGACCGGAAGAGGTACACCAGCACCAAAGAG
GTGCTGGACGCCACCCTGATCCACCAGAGCATCACCGGCCTGTACGAGACACGG
ATCGACCTGTCTCAGCTGGGAGGCGACAAAAGGCCGGCGGCCACGAAAAAGGCC
GGCCAGGCAAAAAAGAAAAAGgaattcGGCAGTGGAGAGGGCAGAGGAAGTCTGC
TAACATGCGGTGACGTCGAGGAGAATCCTGGCCCAatgaccgagtacaagcccacggtgcgcctc
gccacccgcgacgacgtccccagggccgtacgcaccctcgccgccgcgttcgccgactaccccgccacgcgccacaccgtcgatc
cggaccgccacatcgagcgggtcaccgagctgcaagaactcttcctcacgcgcgtcgggctcgacatcggcaaggtgtgggtcgc
ggacgacggcgccgcggtggcggtctggaccacgccggagagcgtcgaagcgggggcggtgttcgccgagatcggcccgcgc
atggccgagttgagcggttcccggctggccgcgcagcaacagatggaaggcctcctggcgccgcaccggcccaaggagcccgcg
tggttcctggccaccgtcggcgtctcgcccgaccaccagggcaagggtctgggcagcgccgtcgtgctccccggagtggaggcgg
ccgagcgcgccggggtgcccgccttcctggagacctccgcgccccacaacctccccttctacgagcggctcggcttcaccgtcacc
gccgacgtcgaggtgcccgaaggaccgcgcacctggtgcatgacccgcaagcccggtgcctgagaattctaaCTAGAGCT
CGCTGATCAGCCTCGACTGTGCCTTCTAGTTGCCAGCCATCTGTTGTTTGCCCCTC
CCCCGTGCCTTCCTTGACCCTGGAAGGTGCCACTCCCACTGTCCTTTCCTAATAAA
ATGAGGAAATTGCATCGCATTGTCTGAGTAGGTGTCATTCTATTCTGGGGGGTGG
GGTGGGGCAGGACAGCAAGGGGGAGGATTGGGAAGAgAATAGCAGGCATGCTG
GGGAgcggccgcaggaacccctagtgatggagttggccactccctctctgcgcgctcgctcgctcactgaggccgggcgacca
aaggtcgcccgacgcccgggctttgcccgggcggcctcagtgagcgagcgagcgcgcagctgcctgcaggggcgcctgatgcg
gtattttctccttacgcatctgtgcggtatttcacaccgcatacgtcaaagcaaccatagtacgcgccctgtagcggcgcattaagcgcg
gcgggtgtggtggttacgcgcagcgtgaccgctacacttgccagcgccctagcgcccgctcctttcgctttcttcccttcctttctcgcca
cgttcgccggctttccccgtcaagctctaaatcgggggctccctttagggttccgatttagtgctttacggcacctcgaccccaaaaaact
tgatttgggtgatggttcacgtagtgggccatcgccctgatagacggtttttcgccctttgacgttggagtccacgttctttaatagtggact
cttgttccaaactggaacaacactcaaccctatctcgggctattcttttgatttataagggattttgccgatttcggcctattggttaaaaaat
gagctgatttaacaaaaatttaacgcgaattttaacaaaatattaacgtttacaattttatggtgcactctcagtacaatctgctctgatgccg
catagttaagccagccccgacacccgccaacacccgctgacgcgccctgacgggcttgtctgctcccggcatccgcttacagacaag
ctgtgaccgtctccgggagctgcatgtgtcagaggttttcaccgtcatcaccgaaacgcgcgagacgaaagggcctcgtgatacgcct

attttataggttaatgtcatgataataatggtttcttagacgtcaggtggcacttttcggggaaatgtgcgcggaacccctatttgtttattttt
ctaaatacattcaaatatgtatccgctcatgagacaataaccctgataaatgcttcaataatattgaaaaaggaagagtatgagtattcaac
atttccgtgtcgcccttattcccttttttgcggcattttgccttcctgtttttgctcacccagaaacgctggtgaaagtaaaagatgctgaaga
tcagttgggtgcacgagtgggttacatcgaactggatctcaacagcggtaagatccttgagagttttcgccccgaagaacgttttccaat
gatgagcacttttaaagttctgctatgtggcgcggtattatcccgtattgacgccgggcaagagcaactcggtcgccgcatacactattct
cagaatgacttggttgagtactcaccagtcacagaaaagcatcttacggatggcatgacagtaagagaattatgcagtgctgccataac
catgagtgataacactgcggccaacttacttctgacaacgatcggaggaccgaaggagctaaccgcttttttgcacaacatgggggat
catgtaactcgccttgatcgttgggaaccggagctgaatgaagccataccaaacgacgagcgtgacaccacgatgcctgtagcaatg
gcaacaacgttgcgcaaactattaactggcgaactacttactctagcttcccggcaacaattaatagactggatggaggcggataaagtt
gcaggaccacttctgcgctcggcccttccggctggctggtttattgctgataaatctggagccggtgagcgtggaagccgcggtatcat
tgcagcactggggccagatggtaagccctcccgtatcgtagttatctacacgacggggagtcaggcaactatggatgaacgaaatag
acagatcgctgagataggtgcctcactgattaagcattggtaactgtcagaccaagtttactcatatatactttagattgatttaaaacttcat
ttttaatttaaaaggatctaggtgaagatcctttttgataatctcatgaccaaaatcccttaacgtgagttttcgttccactgagcgtcagacc
ccgtagaaaagatcaaaggatcttcttgagatcctttttttctgcgcgtaatctgctgcttgcaaacaaaaaaaccaccgctaccagcggt
ggtttgtttgccggatcaagagctaccaactctttttccgaaggtaactggcttcagcagagcgcagataccaaatactgtccttctagtgt
agccgtagttaggccaccacttcaagaactctgtagcaccgcctacatacctcgctctgctaatcctgttaccagtggctgctgccagtg
gcgataagtcgtgtcttaccgggttggactcaagacgatagttaccggataaggcgcagcggtcgggctgaacggggggttcgtgca
cacagcccagcttggagcgaacgacctacaccgaactgagatacctacagcgtgagctatgagaaagcgccacgcttcccgaagg
gagaaaggcggacaggtatccggtaagcggcagggtcggaacaggagagcgcacgagggagcttccagggggaaacgcctggt
atctttatagtcctgtcgggtttcgccacctctgacttgagcgtcgatttttgtgatgctcgtcaggggggcggagcctatggaaaaacgc
cagcaacgcggcctttttacggttcctggccttttgctggccttttgctcacatgt

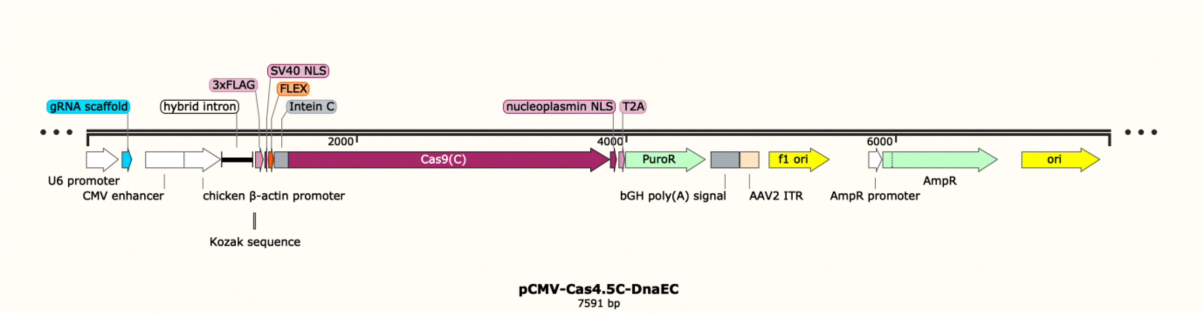


**Nucleic acid Sequence of the construct $_{p}$hCEA-Cas4.5$_{N}$-DnaE$_{N}$ (6917 bp)**

gagggcctatttcccatgattccttcatatttgcatatacgatacaaggctgttagagagataattggaattaatttgactgtaaacacaaag
atattagtacaaaatacgtgacgtagaaagtaataatttcttgggtagtttgcagttttaaaattatgttttaaaatggactatcatatgcttacc
gtaacttgaaagtatttcgatttcttggctttatatatcttGTGGAAAGGACGAAACACCggGTCTTCgaGAAG
ACCTGTTTTAGAGCTAGAAATAGCAAGTTAAAATAAGGCTAGTCCGTTATCAACT
TGAAAAAGTGGCACCGAGTCGGTGcTTTTTTgttttagagctagaaatagcaagttaaaataaggctagtc
cgtTTTTagcgcgtgcgccaattctgcagacaaatggctctagaggtaccGCCCTGGAGAGCATGGGGAGAC
CCGGGACCCTGCTGGGTTTCTCTGTCACAAAGGAAAATAATCCCCCTGGTGTGAC
AGACCCAAGGACAGAACACAGCAGAGGTCAGCACTGGGGAAGACAGGTTGTCC
TCCCAGGGGATGGGGGTCCATCCACCTTGCCGAAAAGATTTGTCTGAGGAACTG
AAAATAGAAGGGAAAAAAGAGGAGGGACAAAAGAGGCAGAAATGAGAGGGGA
GGGGACAGAGGACACCTGAATAAAGACCACACCCATGACCCACGTGATGCTGAG
AAGTACTCCTGCCCTAGGAAGAGACTCAGGGCAGAGGGAGGAAGGACAGCAGA
CCAGACAGTCACAGCAGCCTTGACAAAACGTTCCTGGAACTCAAGCTCTTCTCCA

CAGAGGAGGACAGAGCAGACAGCAGAGACCggagtcgctgcgacgctgccttcgccccgtgccccgc
tccgccgccgcctcgcgccgcccgccccggctctgactgaccgcgttactcccacaggtgagcgggcgggacggcccttctcctcc
gggctgtaattagctgagcaagaggtaagggtttaagggatggttggttggtggggtattaatgtttaattacctggagcacctgcctga
aatcactttttttcaggttGGaccggtgccaccATGGACTATAAGGACCACGACGGAGACTACAAGG
ATCATGATATTGATTACAAAGACGATGACGATAAGATGGCCCCAAAGAAGAAGC
GGAAGGTCGGTATCCACGGAGTCCCAGCAGCCGACAAGAAGTACAGCATCGGCC
TGGACATCGGCACCAACTCTGTGGGCTGGGCCGTGATCACCGACGAGTACAAGG
TGCCCAGCAAGAAATTCAAGGTGCTGGGCAACACCGACCGGCACAGCATCAAGA
AGAACCTGATCGGAGCCCTGCTGTTCGACAGCGGCGAAACAGCCGAGGCCACCC
GGCTGAAGAGAACCGCCAGAAGAAGATACACCAGACGGAAGAACCGGATCTGC
TATCTGCAAGAGATCTTCAGCAACGAGATGGCCAAGGTGGACGACAGCTTCTTC
CACAGACTGGAAGAGTCCTTCCTGGTGGAAGAGGATAAGAAGCACGAGCGGCAC
CCCATCTTCGGCAACATCGTGGACGAGGTGGCCTACCACGAGAAGTACCCCACC
ATCTACCACCTGAGAAAGAAACTGGTGGACAGCACCGACAAGGCCGACCTGCGG
CTGATCTATCTGGCCCTGGCCCACATGATCAAGTTCCGGGGCCACTTCCTGATCG
AGGGCGACCTGAACCCCGACAACAGCGACGTGGACAAGCTGTTCATCCAGCTGG
TGCAGACCTACAACCAGCTGTTCGAGGAAAACCCCATCAACGCCAGCGGCGTGG
ACGCCAAGGCCATCCTGTCTGCCAGACTGAGCAAGAGCAGACGGCTGGAAAATC
TGATCGCCCAGCTGCCCGGCGAGAAGAAGAATGGCCTGTTCGGAAACCTGATTG
CCCTGAGCCTGGGCCTGACCCCCAACTTCAAGAGCAACTTCGACCTGGCCGAGG
ATGCCAAACTGCAGCTGAGCAAGGACACCTACGACGACGACCTGGACAACCTGC
TGGCCCAGATCGGCGACCAGTACGCCGACCTGTTTCTGGCCGCCAAGAACCTGTC
CGACGCCATCCTGCTGAGCGACATCCTGAGAGTGAACACCGAGATCACCAAGGC
CCCCCTGAGCGCCTCTATGATCAAGAGATACGACGAGCACCACCAGGACCTGAC
CCTGCTGAAAGCTCTCGTGCGGCAGCAGCTGCCTGAGAAGTACAAAGAGATTTT
CTTCGACCAGAGCAAGAACGGCTACGCCGGCTACATTGACGGCGGAGCCAGCCA
GGAAGAGTTCTACAAGTTCATCAAGCCCATCCTGGAAAAGATGGACGGCACCGA
GGAACTGCTCGTGAAGCTGAACAGAGAGGACCTGCTGCGGAAGCAGCGGACCTT
CGACAACGGCAGCATCCCCCACCAGATCCACCTGGGAGAGCTGCACGCCATTCT
GCGGCGGCAGGAAGATTTTTACCCATTCCTGAAGGACAACCGGGAAAAGATCGA
GAAGATCCTGACCTTCCGCATCCCCTACTACGTGGGCCCTCTGGCCAGGGGAAAC
AGCAGATTCGCCTGGATGACCAGAAAGAGCGAGGAAACCATCACCCCCTGGAAC
TTCGAGGAAGTGGTGGACAAGGGCGCTTCCGCCCAGAGCTTCATCGAGCGGATG
ACCAACTTCGATAAGAACCTGCCCAACGAGAAGGTGCTGCCCAAGCACAGCCTG
CTGTACGAGTACTTCACCGTGTATAACGAGCTGACCAAAGTGAAATACGTGACC
GAGGGAATGAGAAAGCCCGCCTTCCTGAGCGGCGAGCAGAAAAAGGCCATCGT
GGACCTGCTGTTCAAGACCAACCGGAAAGTGACCGTGAAGCAGCTGAAAGAGGA
CTACTTCAAGAAAATCGAGGGATCCTGTTTAAGCTATGAAACGGAAATATTGAC
AGTAGAATATGGATTATTACCGATTGGTAAAATTGTAGAAAAGCGCATCGAATG
TACTGTTTATAGCGTTGATAATAATGGAAATATTTATACACAACCTGTAGCACAA
TGGCACGATCGCGGAGAACAAGAGGTGTTTGAGTATTGTTTGGAAGATGGTTCA
TTGATTCGGGCAACAAAAGACCATAAGTTTATGACTGTTGATGGTCAAATGTTGC
CAATTGATGAAATATTTGAACGTGAATTGGATTTGATGCGGGTTGATAATTTGCC
GAATgaattcGGCAGTGGAGAGGGCAGAGGAAGTCTGCTAACATGCGGTGACGTCG
AGGAGAATCCTGGCCCAatgaccgagtacaagcccacggtgcgcctcgccacccgcgacgacgtccccagggcc
gtacgcaccctcgccgccgcgttcgccgactaccccgccacgcgccacaccgtcgatccggaccgccacatcgagcgggtcaccg
agctgcaagaactcttcctcacgcgcgtcgggctcgacatcggcaaggtgtgggtcgcggacgacggcgccgcggtggcggtctg
gaccacgccggagagcgtcgaagcgggggcggtgttcgccgagatcggcccgcgcatggccgagttgagcggttcccggctggc
cgcgcagcaacagatggaaggcctcctggcgccgcaccggcccaaggagcccgcgtggttcctggccaccgtcggcgtctcgcc
cgaccaccagggcaagggtctgggcagcgccgtcgtgctccccggagtggaggcggccgagcgcgccggggtgcccgccttcct

ggagacctccgcgccccacaacctccccttctacgagcggctcggcttcaccgtcaccgccgacgtcgaggtgcccgaaggaccg
cgcacctggtgcatgacccgcaagcccggtgcctgagaattctaaCTAGAGCTCGCTGATCAGCCTCGACT
GTGCCTTCTAGTTGCCAGCCATCTGTTGTTTGCCCCTCCCCCGTGCCTTCCTTGAC
CCTGGAAGGTGCCACTCCCACTGTCCTTTCCTAATAAAATGAGGAAATTGCATCG
CATTGTCTGAGTAGGTGTCATTCTATTCTGGGGGGTGGGGTGGGGCAGGACAGC
AAGGGGGAGGATTGGGAAGAgAATAGCAGGCATGCTGGGGAgcggccgcaggaacccta
gtgatggagttggccactccctctctgcgcgctcgctcgctcactgaggccgggcgaccaaaggtcgcccgacgcccgggctttgcc
cgggcggcctcagtgagcgagcgagcgcgcagctgcctgcaggggcgcctgatgcggtattttctccttacgcatctgtgcggtattt
cacaccgcatacgtcaaagcaaccatagtacgcgccctgtagcggcgcattaagcgcggcgggtgtggtggttacgcgcagcgtga
ccgctacacttgccagcgccctagcgcccgctcctttcgctttcttcccttcctttctcgccacgttcgccggctttccccgtcaagctcta
aatcgggggctccctttagggttccgatttagtgctttacggcacctcgaccccaaaaaacttgatttgggtgatggttcacgtagtgggc
catcgccctgatagacggtttttcgccctttgacgttggagtccacgttctttaatagtggactcttgttccaaactggaacaacactcaac
cctatctcgggctattcttttgatttataagggattttgccgatttcggcctattggttaaaaaatgagctgatttaacaaaaatttaacgcgaa
ttttaacaaaatattaacgtttacaattttatggtgcactctcagtacaatctgctctgatgccgcatagttaagccagccccgacacccgcc
aacacccgctgacgcgccctgacgggcttgtctgctcccggcatccgcttacagacaagctgtgaccgtctccgggagctgcatgtgt
cagaggttttcaccgtcatcaccgaaacgcgcgagacgaaagggcctcgtgatacgcctatttttataggttaatgtcatgataataatg
gtttcttagacgtcaggtggcacttttcggggaaatgtgcgcggaacccctatttgtttatttttctaaatacattcaaatatgtatccgctcat
gagacaataaccctgataaatgcttcaataatattgaaaaaggaagagtatgagtattcaacatttccgtgtcgcccttattcccttttttgc
ggcattttgccttcctgtttttgctcacccagaaacgctggtgaaagtaaaagatgctgaagatcagttgggtgcacgagtgggttacatc
gaactggatctcaacagcggtaagatccttgagagttttcgccccgaagaacgttttccaatgatgagcacttttaaagttctgctatgtg
gcgcggtattatcccgtattgacgccgggcaagagcaactcggtcgccgcatacactattctcagaatgacttggttgagtactcacca
gtcacagaaaagcatcttacggatggcatgacagtaagagaattatgcagtgctgccataaccatgagtgataacactgcggccaactt
acttctgacaacgatcggaggaccgaaggagctaaccgcttttttgcacaacatgggggatcatgtaactcgccttgatcgttgggaac
cggagctgaatgaagccataccaaacgacgagcgtgacaccacgatgcctgtagcaatggcaacaacgttgcgcaaactattaactg
gcgaactacttactctagcttcccggcaacaattaatagactggatggaggcggataaagttgcaggaccacttctgcgctcggccctt
ccggctggctggtttattgctgataaatctggagccggtgagcgtggaagccgcggtatcattgcagcactggggccagatggtaagc
cctcccgtatcgtagttatctacacgacggggagtcaggcaactatggatgaacgaaatagacagatcgctgagataggtgcctcact
gattaagcattggtaactgtcagaccaagtttactcatatatactttagattgatttaaaacttcatttttaatttaaaaggatctaggtgaagat
cctttttgataatctcatgaccaaaatcccttaacgtgagttttcgttccactgagcgtcagaccccgtagaaaagatcaaaggatcttcttg
agatcctttttttctgcgcgtaatctgctgcttgcaaacaaaaaaaccaccgctaccagcggtggtttgtttgccggatcaagagctacca
actctttttccgaaggtaactggcttcagcagagcgcagataccaaatactgtccttctagtgtagccgtagttaggccaccacttcaaga
actctgtagcaccgcctacatacctcgctctgctaatcctgttaccagtggctgctgccagtggcgataagtcgtgtcttaccgggttgga
ctcaagacgatagttaccggataaggcgcagcggtcgggctgaacggggggttcgtgcacacagcccagcttggagcgaacgacc
tacaccgaactgagatacctacagcgtgagctatgagaaagcgccacgcttcccgaagggagaaaggcggacaggtatccggtaa
gcggcagggtcggaacaggagagcgcacgagggagcttccagggggaaacgcctggtatctttatagtcctgtcgggtttcgccac
ctctgacttgagcgtcgatttttgtgatgctcgtcaggggggcggagcctatggaaaaacgccagcaacgcggcctttttacggttcctg
gccttttgctggccttttgctcacatgt

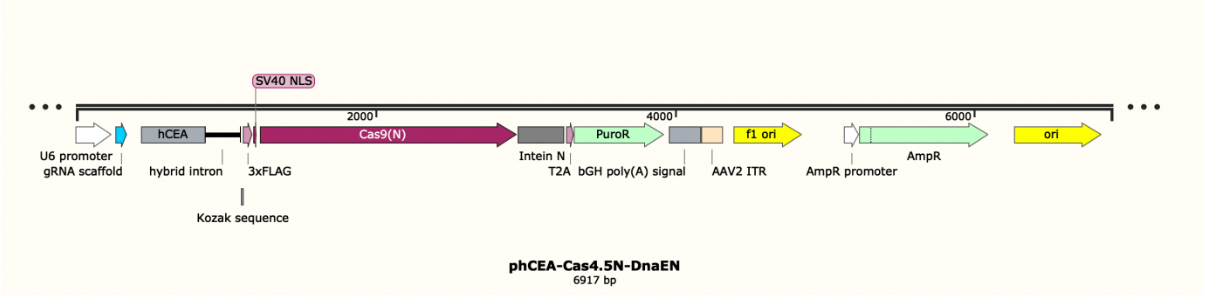

**Nucleic acid Sequence of the construct Twist1BD-Cas4.5$_N$-DnaE$_N$ (6931 bp)**

gagctcatcgatctcgacattgattattgactagttattaatagtaatcaattacggggtcattagttcatagcccatatatggagttccgcgt
tacataacttacggtaaatggcccgcctggctgaccgcccaacgacccccgcccattgacgtcaataatgacgtatgttcccatagtaa
cgccaatagggactttccattgacgtcaatgggtggactatttacggtaaactgcccacttggcagtacatcaagtgtatcatatgccaa
gtacgccccctattgacgtcaatgacggtaaatggcccgcctggcattatgcccagtacatgaccttatgggactttcctacttggcagt
acatctacgtattagtcatcgctattaccatgggtcgaggtgagccccacgttctgcttcactctccccatctcccccccctccccacccc
caattttgtatttatttattttttaattattttgtgcagcgatgggggcggggggggggggggcgcgcgccaggcggggcggggcggg
gcgaggggcggggcggggcgaggcggagaggtgcggcggcagccaatcagagcggcgcgctccgaaagtttccttttatggcg
aggcggcggcggcggcggccctataaaaagcgaagcgcgcggcgggcgggagtcgctgcgttgccttcgccccgtgccccgctc
cgcgccgcctcgcgccgcccgccccggctctgactgaccgcgttactcccacaggtgagcgggcgggacggcccttctcctccgg
gctgtaattagcgcttggtttaatgacggctcgtttcttttctgtggctgcgtgaaagccttaaagggctccgggagggccctttgtgcgg
gggggagcggctcggggggtgcgtgcgtgtgtgtgtgcgtggggagcgccgcgtgcggcccgcgctgcccggcggctgtgagc
gctgcgggcgcggcgcggggctttgtgcgctccgcgtgtgcgcgaggggagcgcggccgggggcggtgccccgcggtgcggg
gggctgcgaggggaacaaaggctgcgtgcggggtgtgtgcgtgggggggtgagcagggggtgtgggcgcggcggtcgggct
gtaacccccccctgcacccccctccccgagttgctgagcacggcccggcttcgggtgcggggctccgtgcggggcgtggcgcggg
gctcgccgtgccgggcggggggtggcggcaggtgggggtgccgggcggggcggggccgcctcgggccggggagggctcgg
gggaggggcgcggcggccccggagcgccggcggctgtcgaggcgcggcgagccgcagccattgccttttatggtaatcgtgcga
gagggcgcagggacttcctttgtcccaaatctggcggagccgaaatctgggaggcgccgccgcaccccctctagcgggcgcgggc
gaagcggtgcggcgccggcaggaaggaaatgggcggggagggccttcgtgcgtcgccgcgccgccgtccccttctccatctcca
gcctcggggctgccgcagggggacggctgccttcgggggggacggggcagggcggggttcggcttctggcgtgtgaccggcgg
ctctagcctctgctaaccatgttcatgccttcttctttttcctacagctcctgggcaacgtgctggttattgtgctgtctcatcattttggcaaat
ctagagccgccatggtgagcaagggcgaggagctgttcaccggggtggtgcccatcctggtcgagctggacggcgacgtaaacgg
ccacaagttcagcgtgtccggcgagggcgagggcgatgccacctacggcaagctgaccctgaagttcatctgcaccaccggcaag
ctgcccgtgccctggcccaccctcgtgaccaccctgacctacggcgtgcagtgcttcagccgctaccccgaccacatgaagcagcac
gacttcttcaagtccgccatgcccgaaggctacgtccaggagcgcaccatcttcttcaaggacgacggcaactacaagacccgcgcc
gaggtgaagttcgagggcgacaccctggtgaaccgcatcgagctgaagggcatcgacttcaaggaggacggcaacatcctggggc
acaagctggagtacaactacaacagccacaacgtctatatcatggccgacaagcagaagaacggcatcaaggtgaacttcaagatcc
gccacaacatcgaggacggcagcgtgcagctcgccgaccactaccagcagaacacccccatcggcgacggccccgtgctgctgc
ccgacaaccactgaggatccgctagcctgcaggtcgacgaattcgatatcggcaagctgaccctgaagttcatctgcaccaccggca
agctgcccgtgccctggcccaccctcgtgaccaccctgacctacggcgtgcagtgcttcagccgctaccccgaccacatgaagcag
cacgacttcttcaagtccgccatgcccgaaggctacgtccaggagcgcaccatcttcttcaaggacgacggcaactacaagacccgc
gccgaggtgaagttcgagggcgacaccctggtgaaccgcatcgagctgaagggcatcgacttcaaggaggacggcaacatcctgg
ggcacaagctggagtacaactacaacagccacaacgtctatatcatggccgacaagcagaagaacggcatcaaggtgaacttcaag
atccgccacaacatcgaggacggcagcgtgcagctcgccgaccactaccagcagaacacccccatcggcgacggccccgtgctg
ctgcccgacaaccactacctgagcacccagtccgccctgagcaaagaccccaacgagaagcgcgatcacatggtcctgctggagtt
cgtgaccgccgccgggatcactctcggcatggacgagctgtacaagtaactcgagactcctcaggtgcaggctgcctatcagaaggt
ggtggctggtgtggccaatgccctggctcacaaataccactgagatctttttccctctgccaaaaattatggggacatcatgaagcccct
tgagcatctgacttctggctaataaaggaaatttattttcattgcaatagtgtgttggaattttttgtgtctctcactcggaaggacatatggg
agggcaaatcatttaaaacatcagaatgagtatttggtttagagtttggcaacatatgcccatatgctggctgccatgaacaaaggttggc
tataaagaggtcatcagtatatgaaacagccccctgctgtccattccttattccatagaaaagccttgacttgaggttagattttttttatatttt
gttttgtgttattttttttctttaacatccctaaaattttccttacatgttttactagccagatttttcctcctctcctgactactcccagtcatagctgt
ccctcttctcttatgaagatccctcgacttaattaaggtacccaattcgccctatagtgagtcgtattacgcgcgctcactggccgtcgtttt
acaacgtcgtgactgggaaaaccctggcgttacccaacttaatcgccttgcagcacatccccctttcgccagctggcgtaatagcgaa
gaggcccgcaccgatcgcccttcccaacagttgcgcagcctgaatggcgaatgggacgcgccctgtagcggcgcattaagcgcgg
cgggtgtggtggttacgcgcagcgtgaccgctacacttgccagcgccctagcgcccgctcctttcgctttcttcccttcctttctcgccac
gttcgccggctttccccgtcaagctctaaatcgggggctccctttagggttccgatttagtgctttacggcacctcgaccccaaaaaactt
gattagggtgatggttcacgtagtgggccatcgccctgatagacggtttttcgccctttgacgttggagtccacgttctttaatagtggact
cttgttccaaactggaacaacactcaaccctatctcggtctattcttttgatttataagggattttgccgatttcggcctattggttaaaaaatg

agctgatttaacaaaaatttaacgcgaattttaacaaaatattaacgcttacaatttaggtggcacttttcggggaaatgtgcgcggaaccc
ctatttgtttatttttctaaatacattcaaatatgtatccgctcatgagacaataaccctgataaatgcttcaataatattgaaaaaggaagagt
atgagtattcaacatttccgtgtcgcccttattcccttttttgcggcattttgccttcctgtttttgctcacccagaaacgctggtgaaagtaaa
agatgctgaagatcagttgggtgcacgagtgggttacatcgaactggatctcaacagcggtaagatccttgagagttttcgccccgaag
aacgttttccaatgatgagcacttttaaagttctgctatgtggcgcggtattatcccgtattgacgccgggcaagagcaactcggtcgcc
gcatacactattctcagaatgacttggttgagtactcaccagtcacagaaaagcatcttacggatggcatgacagtaagagaattatgca
gtgctgccataaccatgagtgataacactgcggccaacttacttctgacaacgatcggaggaccgaaggagctaaccgcttttttgcac
aacatgggggatcatgtaactcgccttgatcgttgggaaccggagctgaatgaagccataccaaacgacgagcgtgacaccacgatg
cctgtagcaatggcaacaacgttgcgcaaactattaactggcgaactacttactctagcttcccggcaacaattaatagactggatggag
gcggataaagttgcaggaccacttctgcgctcggcccttccggctggctggtttattgctgataaatctggagccggtgagcgtgggtc
tcgcggtatcattgcagcactggggccagatggtaagccctcccgtatcgtagttatctacacgacggggagtcaggcaactatggat
gaacgaaatagacagatcgctgagataggtgcctcactgattaagcattggtaactgtcagaccaagtttactcatatatactttagattg
atttaaaacttcatttttaatttaaaaggatctaggtgaagatcctttttgataatctcatgaccaaaatcccttaacgtgagttttcgttccact
gagcgtcagaccccgtagaaaagatcaaaggatcttcttgagatcctttttttctgcgcgtaatctgctgcttgcaaacaaaaaaaccacc
gctaccagcggtggtttgtttgccggatcaagagctaccaactctttttccgaaggtaactggcttcagcagagcgcagataccaaatac
tgtccttctagtgtagccgtagttaggccaccacttcaagaactctgtagcaccgcctacatacctcgctctgctaatcctgttaccagtgg
ctgctgccagtggcgataagtcgtgtcttaccgggttggactcaagacgatagttaccggataaggcgcagcggtcgggctgaacgg
ggggttcgtgcacacagcccagcttggagcgaacgacctacaccgaactgagatacctacagcgtgagctatgagaaagcgccacg
cttcccgaagggagaaaggcggacaggtatccggtaagcggcagggtcggaacaggagagcgcacgagggagcttccaggggg
aaacgcctggtatctttatagtcctgtcgggtttcgccacctctgacttgagcgtcgatttttgtgatgctcgtcaggggggcggagcctat
ggaaaaacgccagcaacgcggcctttttacggttcctggccttttgctggccttttgctcacatgttctttcctgcgttatcccctgattctgt
ggataaccgtattaccgcctttgagtgagctgataccgctcgccgcagccgaacgaccgagcgcagcgagtcagtgagcgaggaa
gcggaagagcgcccaatacgcaaaccgcctctccccgcgcgttggccgattcattaatgcagctggcacgacaggtttcccgactgg
aaagcgggcagtgagcgcaacgcaattaatgtgagttagctcactcattaggcaccccaggctttacactttatgcttccggctcgtatg
ttgtgtggaattgtgagcggataacaatttcacacaggaaacagctatgaccatgattacgccaagcgcgcaattaaccctcactaaag
ggaacaaaagctg

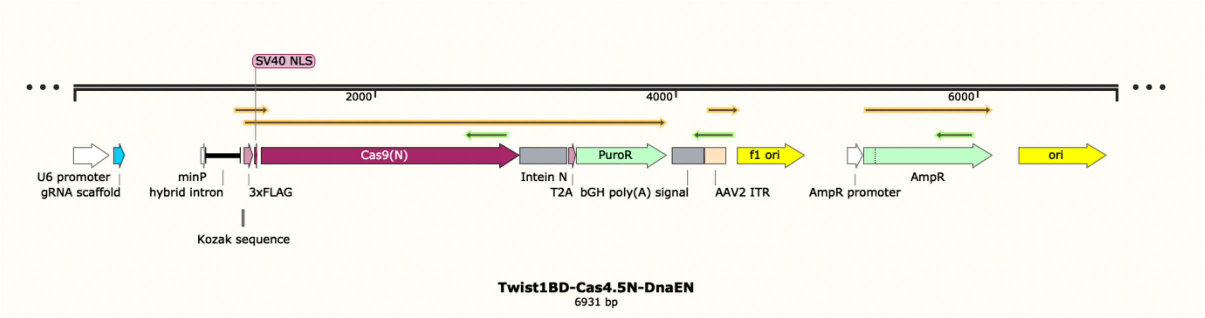


### Nucleic acid Sequence of the construct pCAG-EGxxFP (6383 bp)

gagctcatcgatctcgacattgattattgactagttattaatagtaatcaattacggggtcattagttcatagcccatatatggagttccgcgt
tacataacttacggtaaatggcccgcctggctgaccgcccaacgacccccgcccattgacgtcaataatgacgtatgttcccatagtaa
cgccaatagggactttccattgacgtcaatgggtggactatttacggtaaactgcccacttggcagtacatcaagtgtatcatatgccaa
gtacgccccctattgacgtcaatgacggtaaatggcccgcctggcattatgcccagtacatgaccttatgggactttcctacttggcagt
acatctacgtattagtcatcgctattaccatgggtcgaggtgagccccacgttctgcttcactctccccatctcccccccctccccaccccc
aattttgtatttatttattttttaattattttgtgcagcgatgggggcgggggggggggggggcgcgcgccaggcggggcggggcggg
gcgaggggcggggcggggcgaggcggagaggtgcggcggcagccaatcagagcggcgcgctccgaaagtttccttttatggcg
aggcggcggcggcggcggccctataaaaagcgaagcgcgcggcgggcgggagtcgctgcgttgccttcgccccgtgccccgctc
cgcgccgcctcgcgccgcccgccccggctctgactgaccgcgttactcccacaggtgagcgggcgggacggcccttctcctccgg

gctgtaattagcgcttggtttaatgacggctcgtttcttttctgtggctgcgtgaaagccttaaagggctccgggagggccctttgtgcgg
gggggagcggctcgggggtgcgtgcgtgtgtgtgcgtggggagcgccgcgtgcggcccgcgctgcccggcggctgtgagc
gctgcgggcgcggcgcggggctttgtgcgctccgcgtgtgcgcgaggggagcgcggccgggggcggtgccccgcggtgcggg
ggggctgcgaggggaacaaaggctgcgtgcggggtgtgtgcgtgggggggtgagcagggggtgtgggcgcggcggtcgggct
gtaacccccccctgcacccccctccccgagttgctgagcacggcccggcttcgggtgcggggctccgtgcggggcgtggcgcggg
gctcgccgtgccgggcggggggtggcggcaggtgggggtgccgggcggggcggggccgcctcgggccggggagggctcgg
gggaggggcgcggcggccccggagcgccggcggctgtcgaggcgcggcgagccgcagccattgccttttatggtaatcgtgcga
gagggcgcagggacttcctttgtcccaaatctggcggagccgaaatctgggaggcgccgccgcaccccctctagcgggcgcgggc
gaagcggtgcggcgccggcaggaaggaaatgggcggggagggccttcgtgcgtcgccgcgccgccgtccccttctccatctcca
gcctcggggctgccgcaggggacggctgccttcgggggggacggggcagggcggggttcggcttctggcgtgtgaccggcgg
ctctagcctctgctaaccatgttcatgccttcttctttttcctacagctcctgggcaacgtgctggttattgtgctgtctcatcattttggcaaat
ctagagccgccatggtgagcaagggcgaggagctgttcaccggggtggtgcccatcctggtcgagctggacggcgacgtaaacgg
ccacaagttcagcgtgtccggcgagggcgagggcgatgccacctacggcaagctgaccctgaagttcatctgcaccaccggcaag
ctgcccgtgccctggcccaccctcgtgaccaccctgacctacggcgtgcagtgcttcagccgctaccccgaccacatgaagcagcac
gacttcttcaagtccgccatgcccgaaggctacgtccaggagcgcaccatcttcttcaaggacgacggcaactacaagacccgcgcc
gaggtgaagttcgagggcgacaccctggtgaaccgcatcgagctgaagggcatcgacttcaaggaggacggcaacatcctggggc
acaagctggagtacaactacaacagccacaacgtctatatcatggccgacaagcagaagaacggcatcaaggtgaacttcaagatcc
gccacaacatcgaggacggcagcgtgcagctcgccgaccactaccagcagaacacccccatcggcgacggccccgtgctgctgc
ccgacaaccactgaggatccgctagcctgcaggtcgacgaattcgatatcggcaagctgaccctgaagttcatctgcaccaccggca
agctgcccgtgccctggcccaccctcgtgaccaccctgacctacggcgtgcagtgcttcagccgctaccccgaccacatgaagcag
cacgacttcttcaagtccgccatgcccgaaggctacgtccaggagcgcaccatcttcttcaaggacgacggcaactacaagacccgc
gccgaggtgaagttcgagggcgacaccctggtgaaccgcatcgagctgaagggcatcgacttcaaggaggacggcaacatcctgg
ggcacaagctggagtacaactacaacagccacaacgtctatatcatggccgacaagcagaagaacggcatcaaggtgaacttcaag
atccgccacaacatcgaggacggcagcgtgcagctcgccgaccactaccagcagaacacccccatcggcgacggccccgtgctg
ctgcccgacaaccactacctgagcacccagtccgccctgagcaaagaccccaacgagaagcgcgatcacatggtcctgctggagtt
cgtgaccgccgccgggatcactctcggcatggacgagctgtacaagtaactcgagactcctcaggtgcaggctgcctatcagaaggt
ggtggctggtgtggccaatgccctggctcacaaataccactgagatctttttccctctgccaaaaattatggggacatcatgaagcccct
tgagcatctgacttctggctaataaaggaaatttattttcattgcaatagtgtgttggaattttttgtgtctctcactcggaaggacatatggg
agggcaaatcatttaaaacatcagaatgagtatttggtttagagtttggcaacatatgcccatatgctggctgccatgaacaaaggttggc
tataaagaggtcatcagtatatgaaacagccccctgctgtccattccttattccatagaaaagccttgacttgaggttagattttttttatatttt
gttttgtgttatttttttctttaacatccctaaaattttccttacatgttttactagccagatttttcctcctctcctgactactcccagtcatagctgt
ccctcttctcttatgaagatccctcgacttaattaaggtacccaattcgccctatagtgagtcgtattacgcgcgctcactggccgtcgtttt
acaacgtcgtgactgggaaaaccctggcgttacccaacttaatcgccttgcagcacatccccctttcgccagctggcgtaatagcgaa
gaggcccgcaccgatcgcccttcccaacagttgcgcagcctgaatggcgaatgggacgcgccctgtagcggcgcattaagcgcgg
cgggtgtggtggttacgcgcagcgtgaccgctacacttgccagcgccctagcgcccgctcctttcgctttcttcccttcctttctcgccac
gttcgccggctttccccgtcaagctctaaatcgggggctccctttagggttccgatttagtgctttacggcacctcgaccccaaaaaactt
gattagggtgatggttcacgtagtgggccatcgccctgatagacggtttttcgccctttgacgttggagtccacgttctttaatagtggact
cttgttccaaactggaacaacactcaaccctatctcggtctattcttttgatttataagggattttgccgatttcggcctattggttaaaaaatg
agctgatttaacaaaaatttaacgcgaattttaacaaaatattaacgcttacaatttaggtggcacttttcggggaaatgtgcgcggaaccc
ctatttgtttatttttctaaatacattcaaatatgtatccgctcatgagacaataaccctgataaatgcttcaataatattgaaaaaggaagagt
atgagtattcaacatttccgtgtcgcccttattcccttttttgcggcattttgccttcctgtttttgctcacccagaaacgctggtgaaagtaaa
agatgctgaagatcagttgggtgcacgagtgggttacatcgaactggatctcaacagcggtaagatccttgagagttttcgccccgaag
aacgttttccaatgatgagcacttttaaagttctgctatgtggcgcggtattatcccgtattgacgccgggcaagagcaactcggtcgcc
gcatacactattctcagaatgacttggttgagtactcaccagtcacagaaaagcatcttacggatggcatgacagtaagagaattatgca
gtgctgccataaccatgagtgataacactgcggccaacttacttctgacaacgatcggaggaccgaaggagctaaccgcttttttgcac
aacatgggggatcatgtaactcgccttgatcgttgggaaccggagctgaatgaagccataccaaacgacgagcgtgacaccacgatg
cctgtagcaatggcaacaacgttgcgcaaactattaactggcgaactacttactctagcttcccggcaacaattaatagactggatggag
gcggataaagttgcaggaccacttctgcgctcggcccttccggctggctggtttattgctgataaatctggagccggtgagcgtgggtc
tcgcggtatcattgcagcactggggccagatggtaagccctcccgtatcgtagttatctacacgacggggagtcaggcaactatggat
gaacgaaatagacagatcgctgagataggtgcctcactgattaagcattggtaactgtcagaccaagtttactcatatatactttagattg

atttaaaacttcatttttaatttaaaaggatctaggtgaagatcctttttgataatctcatgaccaaaatcccttaacgtgagttttcgttccact
gagcgtcagaccccgtagaaaagatcaaaggatcttcttgagatcctttttttctgcgcgtaatctgctgcttgcaaacaaaaaaaccacc
gctaccagcggtggtttgtttgccggatcaagagctaccaactctttttccgaaggtaactggcttcagcagagcgcagataccaaatac
tgtccttctagtgtagccgtagttaggccaccacttcaagaactctgtagcaccgcctacatacctcgctctgctaatcctgttaccagtgg
ctgctgccagtggcgataagtcgtgtcttaccgggttggactcaagacgatagttaccggataaggcgcagcggtcgggctgaacgg
ggggttcgtgcacacagcccagcttggagcgaacgacctacaccgaactgagatacctacagcgtgagctatgagaaagcgccacg
cttcccgaagggagaaaggcggacaggtatccggtaagcggcagggtcggaacaggagagcgcacgagggagcttccaggggg
aaacgcctggtatctttatagtcctgtcgggtttcgccacctctgacttgagcgtcgatttttgtgatgctcgtcaggggggcggagcctat
ggaaaaacgccagcaacgcggcctttttacggttcctggccttttgctggccttttgctcacatgttctttcctgcgttatcccctgattctgt
ggataaccgtattaccgcctttgagtgagctgataccgctcgccgcagccgaacgaccgagcgcagcgagtcagtgagcgaggaa
gcggaagagcgcccaatacgcaaaccgcctctccccgcgcgttggccgattcattaatgcagctggcacgacaggtttcccgactgg
aaagcgggcagtgagcgcaacgcaattaatgtgagttagctcactcattaggcaccccaggctttacactttatgcttccggctcgtatg
ttgtgtggaattgtgagcggataacaatttcacacaggaaacagctatgaccatgattacgccaagcgcgcaattaaccctcactaaag
ggaacaaaagctg

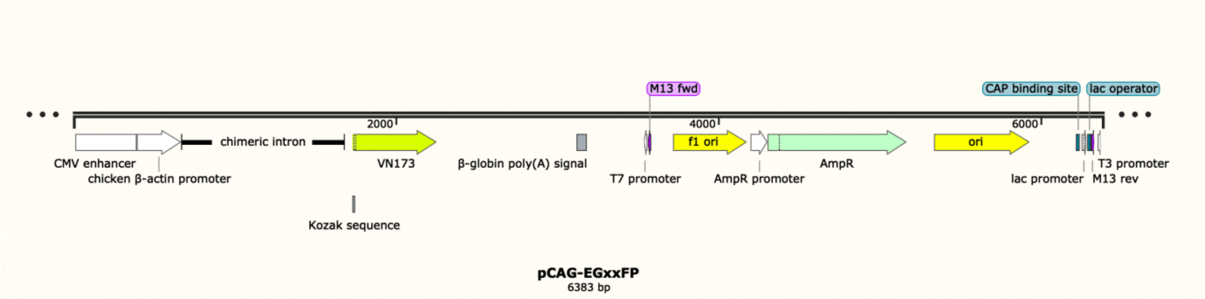


**Nucleic acid Sequence of the construct Syncytin1-P2A-mCherry (6441 bp)**

TAGTTATTAATAGTAATCAATTACGGGGTCATTAGTTCATAGCCCATATATGGAG
TTCCGCGTTACATAACTTACGGTAAATGGCCCGCCTGGCTGACCGCCCAACGACC
CCCGCCCATTGACGTCAATAATGACGTATGTTCCCATAGTAACGCCAATAGGGAC
TTTCCATTGACGTCAATGGGTGGAGTATTTACGGTAAACTGCCCACTTGGCAGTA
CATCAAGTGTATCATATGCCAAGTACGCCCCCTATTGACGTCAATGACGGTAAAT
GGCCCGCCTGGCATTATGCCCAGTACATGACCTTATGGGACTTTCCTACTTGGCA
GTACATCTACGTATTAGTCATCGCTATTACCATGGTGATGCGGTTTTGGCAGTAC
ATCAATGGGCGTGGATAGCGGTTTGACTCACGGGGATTTCCAAGTCTCCACCCCA
TTGACGTCAATGGGAGTTTGTTTTGGCACCAAAATCAACGGGACTTTCCAAAATG
TCGTAACAACTCCGCCCCATTGACGCAAATGGGCGGTAGGCGTGTACGGTGGGA
GGTCTATATAAGCAGAGCTGGTTTAGTGAACCGTCAGATCCCGTCTCcgtacgATGG
CCCTGCCTTACCACATCTTTCTGTTCACCGTGCTGCTGCCCAGCTTCACCCTTACA
GCTCCTCCACCTTGCCGGTGCATGACAAGCAGCAGCCCCTACCAAGAGTTCCTGT
GGCGAATGCAGAGGCCCGGCAATATCGACGCCCCTAGCTACAGAAGCCTGAGCA
AGGGCACCCCTACCTTCACAGCCCACACACACATGCCCCGGAACTGCTACCATA
GCGCCACACTGTGTATGCACGCCAACACACACTACTGGACCGGCAAGATGATCA
ACCCTAGCTGTCCTGGCGGCCTGGGCGTGACAGTGTGTTGGACCTACTTTACCCA
GACCGGCATGTCCGATGGTGGCGGAGTTCAAGACCAGGCCAGAGAAAAGCACGT
GAAAGAAGTGATCAGCCAGCTGACCAGAGTGCACGGCACAAGCAGCCCTTACAA
AGGCCTGGACCTGTCCAAGCTGCACGAGACACTGAGAACCCACACACGGCTGGT
GTCCCTGTTCAACACCACACTGACAGGCCTGCACGAGGTGTCCGCTCAGAACCCT
ACCAACTGCTGGATCTGCCTGCCTCTGAACTTCAGACCCTACGTGTCAATCCCCG

TGCCTGAGCAGTGGAACAACTTCAGCACCGAGATCAATACCACCAGCGTGCTCG
TGGGCCCTCTGGTGTCCAATCTGGAAATCACCCACACCAGCAACCTGACCTGCGT
GAAGTTCTCCAACACCACCTACACCACCAACAGCCAGTGCATCAGATGGGTCAC
CCCTCCAACACAGATCGTGTGTCTGCCCAGCGGCATCTTCTTCGTGTGTGGCACA
AGCGCCTACCGGTGCCTGAATGGCAGCAGCGAGTCTATGTGCTTCCTGAGCTTCC
TGGTGCCTCCTATGACCATCTACACCGAGCAGGACCTGTACAGCTACGTGATCAG
CAAGCCCAGAAACAAGCGGGTGCCCATCCTGCCTTTTGTGATTGGAGCTGGCGTG
CTGGGAGCCCTCGGAACTGGAATCGGAGGCATCACCACCAGCACACAGTTCTAC
TACAAGCTGAGCCAAGAGCTGAACGGCGACATGGAAAGAGTGGCCGACAGCCT
GGTCACACTGCAGGACCAGCTGAATTCTCTGGCTGCCGTGGTGCTGCAGAATCGG
AGAGCACTCGATCTGCTGACAGCCGAGAGAGGCGGCACCTGTCTGTTTCTCGGC
GAGGAATGCTGCTACTACGTGAACCAGAGCGGCATCGTGACCGAGAAAGTGAAA
GAGATCCGGGACAGAATCCAGCGGAGAGCCGAGGAACTGAGAAACACAGGACC
TTGGGGCCTGCTGAGCCAGTGGATGCCTTGGATCCTGCCATTTCTGGGACCTCTG
GCCGCCATCATTCTGCTGCTGCTGTTCGGCCCCTGCATCTTCAACCTGCTGGTCAA
CTTCGTGTCCAGCCGGATCGAGGCCGTGAAGCTGCAGATGGAACCCAAGATGCA
GAGCAAGACCAAGATCTACAGACGGCCCCTGGACAGACCCGCCTCTCCTAGATC
CGACGTGAACGACATCAAGGGCACACCTCCAGAGGAAATCAGCGCCGCTCAGCC
TCTGCTGAGGCCTAATTCTGCCGGAAGCTCTGCGCGCGAGACGGGAAGCgcgcgcG
AGACGGGAAGCGGAGCTACTAACTTCAGCCTGCTGAAGCAGGCTGGCGACGTGG
AGGAGAACCCTGGACCTGGTCTCcggccgATGGTGAGCAAGGGCGAGGAGGATAA
CATGGCCATCATCAAGGAGTTCATGCGCTTCAAGGTGCACATGGAGGGCTCCGT
GAACGGCCACGAGTTCGAGATCGAGGGCGAGGGCGAGGGCCGCCCCTACGAGG
GCACCCAGACCGCCAAGCTGAAGGTGACCAAGGGTGGCCCCCTGCCCTTCGCCT
GGGACATCCTGTCCCCTCAGTTCATGTACGGCTCCAAGGCCTACGTGAAGCACCC
CGCCGACATCCCCGACTACTTGAAGCTGTCCTTCCCCGAGGGCTTCAAGTGGGAG
CGCGTGATGAACTTCGAGGACGGCGGCGTGGTGACCGTGACCCAGGACTCCTCC
CTGCAGGACGGCGAGTTCATCTACAAGGTGAAGCTGCGCGGCACCAACTTCCCC
TCCGACGGCCCCGTAATGCAGAAGAAGACCATGGGCTGGGAGGCCTCCTCCGAG
CGGATGTACCCCGAGGACGGCGCCCTGAAGGGCGAGATCAAGCAGAGGCTGAA
GCTGAAGGACGGCGGCCACTACGACGCTGAGGTCAAGACCACCTACAAGGCCAA
GAAGCCCGTGCAGCTGCCCGGCGCCTACAACGTCAACATCAAGTTGGACATCAC
CTCCCACAACGAGGACTACACCATCGTGGAACAGTACGAACGCGCCGAGGGCCG
CCACTCCACCGGCGGCATGGACGAGCTGTACAAGTCCGGAAACTAGTCTCAGAT
CTCGAGCTCAAGCTTCGAATTCTGCAGTCGACGGTACCGCGGGCCCGGGATCCAC
CGGATCTAGATAACTGATCATAATCAGCCATACCACATTTGTAGAGGTTTTACTT
GCTTTAAAAAACCTCCCACACCTCCCCCTGAACCTGAAACATAAAATGAATGCA
ATTGTTGTTGTTAACTTGTTTATTGCAGCTTATAATGGTTACAAATAAAGCAATA
GCATCACAAATTTCACAAATAAAGCATTTTTTTCACTGCATTCTAGTTGTGGTTTG
TCCAAACTCATCAATGTATCTTAACGCGTAAATTGTAAGCGTTAATATTTTGTTA
AAATTCGCGTTAAATTTTTGTTAAATCAGCTCATTTTTTAACCAATAGGCCGAAA
TCGGCAAAATCCCTTATAAATCAAAAGAATAGACCGAGATAGGGTTGAGTGTTG
TTCCAGTTTGGAACAAGAGTCCACTATTAAAGAACGTGGACTCCAACGTCAAAG
GGCGAAAAACCGTCTATCAGGGCGATGGCCCACTACGTGAACCATCACCCTAAT
CAAGTTTTTTGGGGTCGAGGTGCCGTAAAGCACTAAATCGGAACCCTAAAGGGA
GCCCCCGATTTAGAGCTTGACGGGGAAAGCCGGCGAACGTGGCGAGAAAGGAA
GGGAAGAAAGCGAAAGGAGCGGGCGCTAGGGCGCTGGCAAGTGTAGCGGTCAC
GCTGCGCGTAACCACCACACCCGCCGCGCTTAATGCGCCGCTACAGGGCGCGTC
AGGTGGCACTTTTCGGGGAAATGTGCGCGGAACCCCTATTTGTTTATTTTTCTAA
ATACATTCAAATATGTATCCGCTCATGAGACAATAACCCTGATAAATGCTTCAAT

AATATTGAAAAAGGAAGAGTCCTGAGGCGGAAAGAACCAGCTGTGGAATGTGTG
TCAGTTAGGGTGTGGAAAGTCCCCAGGCTCCCCAGCAGGCAGAAGTATGCAAAG
CATGCATCTCAATTAGTCAGCAACCAGGTGTGGAAAGTCCCCAGGCTCCCCAGC
AGGCAGAAGTATGCAAAGCATGCATCTCAATTAGTCAGCAACCATAGTCCCGCC
CCTAACTCCGCCCATCCCGCCCCTAACTCCGCCCAGTTCCGCCCATTCTCCGCCCC
ATGGCTGACTAATTTTTTTTATTTATGCAGAGGCCGAGGCCGCCTCGGCCTCTGA
GCTATTCCAGAAGTAGTGAGGAGGCTTTTTTGGAGGCCTAGGCTTTTGCAAAGAT
CGATCAAGAGACAGGATGAGGATCGTTTCGCATGATTGAACAAGATGGATTGCA
CGCAGGTTCTCCGGCCGCTTGGGTGGAGAGGCTATTCGGCTATGACTGGGCACA
ACAGACAATCGGCTGCTCTGATGCCGCCGTGTTCCGGCTGTCAGCGCAGGGGCG
CCCGGTTCTTTTTGTCAAGACCGACCTGTCCGGTGCCCTGAATGAACTGCAAGAC
GAGGCAGCGCGGCTATCGTGGCTGGCCACGACGGGCGTTCCTTGCGCAGCTGTG
CTCGACGTTGTCACTGAAGCGGGAAGGGACTGGCTGCTATTGGGCGAAGTGCCG
GGGCAGGATCTCCTGTCATCTCACCTTGCTCCTGCCGAGAAAGTATCCATCATGG
CTGATGCAATGCGGCGGCTGCATACGCTTGATCCGGCTACCTGCCCATTCGACCA
CCAAGCGAAACATCGCATCGAGCGAGCACGTACTCGGATGGAAGCCGGTCTTGT
CGATCAGGATGATCTGGACGAAGAGCATCAGGGGCTCGCGCCAGCCGAACTGTT
CGCCAGGCTCAAGGCGAGCATGCCCGACGGCGAGGATCTCGTCGTGACCCATGG
CGATGCCTGCTTGCCGAATATCATGGTGGAAAATGGCCGCTTTTCTGGATTCATC
GACTGTGGCCGGCTGGGTGTGGCGGACCGCTATCAGGACATAGCGTTGGCTACC
CGTGATATTGCTGAAGAGCTTGGCGGCGAATGGGCTGACCGCTTCCTCGTGCTTT
ACGGTATCGCCGCTCCCGATTCGCAGCGCATCGCCTTCTATCGCCTTCTTGACGA
GTTCTTCtgaGCGGGACTCTGGGGTTCGAAATGACCGACCAAGCGACGCCCAACCT
GCCATCACGAGATTTCGATTCCACCGCCGCCTTCTATGAAAGGTTGGGCTTCGGA
ATCGTTTTCCGGGACGCCGGCTGGATGATCCTCCAGCGCGGGGATCTCATGCTGG
AGTTCTTCGCCCACCCTAGGGGGAGGCTAACTGAAACACGGAAGGAGACAATAC
CGGAAGGAACCCGCGCTATGACGGCAATAAAAAGACAGAATAAAACGCACGGT
GTTGGGTCGTTTGTTCATAAACGCGGGGTTCGGTCCCAGGGCTGGCACTCTGTCG
ATACCCCACCGAGACCCCATTGGGGCCAATACGCCCGCGTTTCTTCCTTTTCCCC
ACCCCACCCCCCAAGTTCGGGTGAAGGCCCAGGGCTCGCAGCCAACGTCGGGGC
GGCAGGCCCTGCCATAGCCTCAGGTTACTCATATATACTTTAGATTGATTTAAAA
CTTCATTTTTAATTTAAAAGGATCTAGGTGAAGATCCTTTTTGATAATCTCATGAC
CAAAATCCCTTAACGTGAGTTTTCGTTCCACTGAGCGTCAGACCCCGTAGAAAAG
ATCAAAGGATCTTCTTGAGATCCTTTTTTTCTGCGCGTAATCTGCTGCTTGCAAAC
AAAAAAACCACCGCTACCAGCGGTGGTTTGTTTGCCGGATCAAGAGCTACCAAC
TCTTTTTCCGAAGGTAACTGGCTTCAGCAGAGCGCAGATACCAAATACTGTCCTT
CTAGTGTAGCCGTAGTTAGGCCACCACTTCAAGAACTCTGTAGCACCGCCTACAT
ACCTCGCTCTGCTAATCCTGTTACCAGTGGCTGCTGCCAGTGGCGATAAGTCGTG
TCTTACCGGGTTGGACTCAAGACGATAGTTACCGGATAAGGCGCAGCGGTCGGG
CTGAACGGGGGGTTCGTGCACACAGCCCAGCTTGGAGCGAACGACCTACACCGA
ACTGAGATACCTACAGCGTGAGCTATGAGAAAGCGCCACGCTTCCCGAAGGGAG
AAAGGCGGACAGGTATCCGGTAAGCGGCAGGGTCGGAACAGGAGAGCGCACGA
GGGAGCTTCCAGGGGGAAACGCCTGGTATCTTTATAGTCCTGTCGGGTTTCGCCA
CCTCTGACTTGAGCGTCGATTTTTGTGATGCTCGTCAGGGGGGCGGAGCCTATGG
AAAAACGCCAGCAACGCGGCCTTTTTACGGTTCCTGGCCTTTTGCTGGCCTTTTG
CTCACATGTTCTTTCCTGCGTTATCCCCTGATTCTGTGGATAACCGTATTACCGCC
ATGCAT

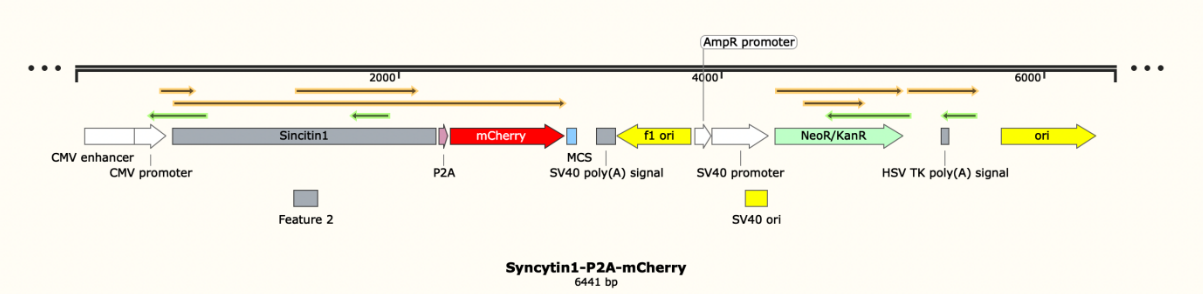
AmpR promoter
2000
4000
6000
Sincitin1
mCherry
f1 ori
NeoR/KanR
ori
CMV enhancer
CMV promoter
P2A
MCS
SV40 poly(A) signal
SV40 promoter
HSV TK poly(A) signal
Feature 2
SV40 ori
Syncytin1-P2A-mCherry
6441 bp
